# Cell division drives DNA methylation loss in late-replicating domains in primary human cells

**DOI:** 10.1101/2022.04.14.485664

**Authors:** Jamie L. Endicott, Paula A. Nolte, Hui Shen, Peter W. Laird

## Abstract

DNA methylation undergoes dramatic age-related changes, first described more than four decades ago^1–4^. Loss of DNA methylation at late-replicating regions of the genome attached to the nuclear lamina advances with age in normal tissues, and is further exacerbated in cancer^5–7^. We present here the first experimental evidence that this DNA hypomethylation is directly driven by proliferation-associated DNA replication. Loss of DNA methylation at low-density CpGs in A:T-rich, <u>p</u>artially <u>m</u>ethylated <u>d</u>omains (PMD solo-WCGWs), tracks cumulative population doublings in primary cell culture. Cell cycle deceleration resulted in a proportional decrease in the rate of DNA hypomethylation. Blocking DNA replication via Mitomycin C treatment halted methylation loss. Loss of methylation continued unabated after TERT immortalization until finally reaching a severely hypomethylated equilibrium. Ambient oxygen culture conditions increased the rate of methylation loss compared to low-oxygen conditions, suggesting that some methylation loss may occur during unscheduled, oxidative damage repair-associated DNA synthesis. Finally, we present and validate a model to estimate the relative cumulative replicative histories of human cells, which we call “RepliTali” (<u>Repli</u>cation <u>T</u>imes <u>A</u>ccumulated in <u>Li</u>fetime).

---

Age-associated DNA hypomethylation is associated with several intertwined spatio-temporal features. DNA methylation loss occurs primarily within PMDs, which largely coincide with late replication timing domains^8,9^, are enriched in higher order chromatin compartment B^10^, and tend to be associated with the nuclear lamina^7^. Cancer-associated DNA methylation loss is accompanied by changes in replication timing and 3D genome organization^11^. Replicative senescence alters 3D genome compartmentalization^12–14^. Replication timing, altered in both cancer and aging-associated diseases including progeria^15–17^, is purported to maintain the epigenome^18,19^, although this relationship may be bidirectional^20^.

Epigenetic ‘clocks’ — models trained upon large DNA methylation datasets to predict either chronological age^21–23^ or features of biological aging^24,25^ — have emerged as powerful tools in aging research in recent years, facilitated by the affordability of DNA methylation microarrays and the subsequent availability of increasingly large publicly available datasets. DNA methylation clocks have far out-performed other metrics of biological age, such as telomere length and transcriptional signatures. Although much focus is on the epigenetic age acceleration that is observed with a multitude of diseases^25,26^, and the slowing or reversal of epigenetic age^27^, recent clock iterations have the intriguing ability to estimate chronological age across mammalian species^28,29^, likely detecting conserved features of aging. Although there have been recent attempts to retroactively classify underlying clock mechanisms^30^, a major limitation to the interpretation of clock results is the lack of understanding of what drives the methylation behaviors of each clock’s CpGs. Whether the age-associated changes in DNA methylation actively contribute to aging, or are merely passenger events, remains largely unknown.

By their nature, chronological methylation clocks are not mitotic clocks. The various tissues within an organism have the same chronological age, but are comprised of cell types with different proliferation rates and replicative histories^31^. DNA methylation clocks calibrated to organismal age therefore need to be impervious to cell type composition differences. This eliminates DNA methylation changes that directly reflect ongoing or past cell division from most epigenetic clocks trained to chronological age using multiple tissues. The process of cell division requires the passage of chronological time, but the two can be unlinked since time can pass without cell division, such as in post-mitotic cells.

It is important to distinguish between replicative history and proliferation rate. Replicative history refers to the cumulative number of cell divisions within a single cell’s lineage. Proliferation rate refers to the number of divisions per unit of time, usually as a current, ongoing measure. In the greater context of biological aging, three of nine ‘Hallmarks of Aging’ are attributable, in great part, to cumulative cell divisions: telomere attrition, stem cell exhaustion, and cellular senescence^4^. Therefore, replicative history is closely tied to biological age and thus an important feature to measure independent of biological measures of chronological time. In light of this, the term ‘clock’ is a misnomer for estimates of cumulative cell divisions. A ‘counter’, ‘enumerator’ or ‘tally’ would more accurately capture the nature of cell division. However, the term ‘epigenetic mitotic clock’ has become cemented into the existing literature for proposed DNA methylation-based measures of cell division^30,32^.

We have previously identified a hypomethylation-prone sequence signature, PMD solo-WCGW, representing PMD CpG dinucleotides immediately flanked by an adenine or thymine (‘W’) and located at least 35bp away from the nearest CpG (‘solo’)^33^ (**Fig. 1b**). PMD solo-WCGW hypomethylation appears to correspond to the approximate replicative history of various tissue types and malignancies, and we hypothesized that this could be attributed to incomplete maintenance methylation at each cell division^6^. Subsequent analyses by other groups confirmed that methylation at PMD solo-WCGWs is indeed maintained poorly relative to other sequence contexts^34^. However, there has been no direct experimental evidence to establish a causal or mechanistic link between replicative history and PMD hypomethylation, and this interpretation has been challenged by others in the field^35,36^.

**Fig. 1.**
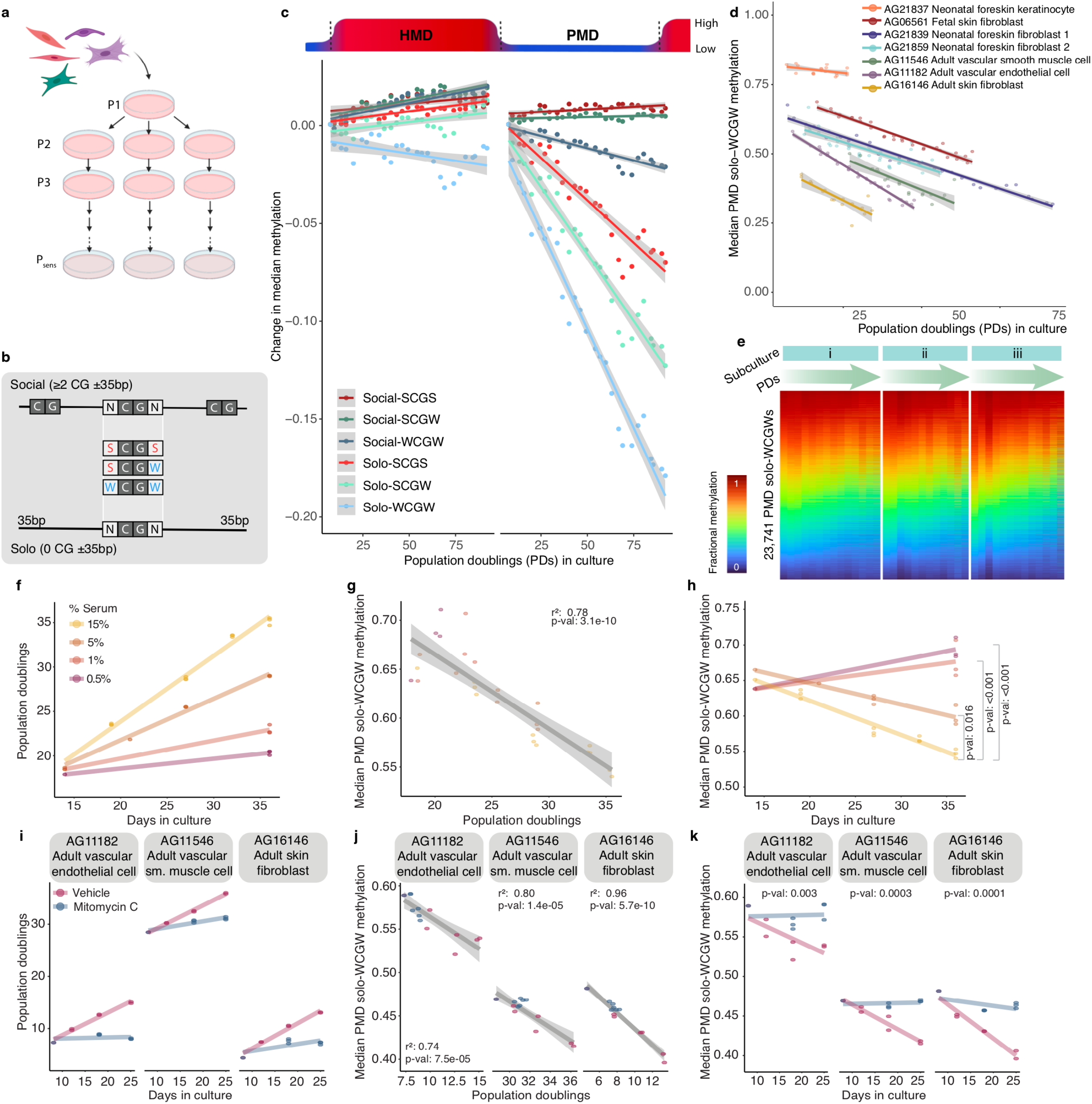
Methylation loss at PMD solo-WCGWs is driven by proliferation-associated DNA replication. **a**, Schematic illustration of primary cells cultured through replicative senescence. **b**, Illustration of immediate (≤35bp) CpG contexts investigated in this study. **c**, Fractional methylation change per population doubling (PD) for neonatal foreskin fibroblast AG21839 at select CpG contexts, within and outside of common partially methylated domain (PMD) boundaries. HMD: highly methylated domains. CpGs within CGIs were excluded. Solid lines depict linear regression with gray shading depicting 95% confidence interval. **d**, Median fractional methylation of multiple primary cell lines plotted against PDs achieved during this study. **e**, DNA methylation heatmap of PMD solo-WCGWs during primary cell culture of neonatal foreskin fibroblast AG21859. Heatmap is separated by parallel subculture, with samples ordered from passage 1 through replicative senescence. **f-h**, Primary cell lines (n=3) transiently treated with DNA crosslinking agent Mitomycin C for 3 hours, followed by drug-free culture for 25 days, resulting in the inhibition of DNA synthesis and subsequent growth arrest, have stable PMD solo-WCGW methylation. **(f)**; PMD solo-WCGW methylation is tightly correlated to PDs **(g)**, independently of time **(h). i**-**k**, Primary fetal skin fibroblast (AG06561) grown in media containing different % v/v fetal bovine serum loses PMD solo-WCGW methylation as a function of mitotic rate. Solid lines depict linear regression with gray shading depicting 95% confidence interval. Statistical comparisons for panels **h** and **k** were performed using mixed effects modeling; p-values adjusted for multiple comparisons presented in panel **k**.

## Context-dependent methylation change in response to cell divisions

We used serial primary human cell cultures to closely track the *in vitro* replication of cell populations. Primary human cell lines (n=7, **Supplementary Table 1**) were obtained from the NIA Aging Cell Culture Repository Apparently Healthy Collection, at the Coriell Institute for Medical Research, and cultured under recommended conditions with multiple parallel subcultures originating from the same initiating cells (**Fig. 1a**) through replicative senescence, tracking cumulative cell divisions (population doublings, PDs) at each passage (methods). At each passaging, a fraction of cells was retained for DNA methylation analysis using the Infinium MethylationEPIC array (Illumina).

Analysis of DNA methylation revealed divergent behavior between non-CGI CpGs within different contexts: CpGs located in PMDs progressively lost methylation, and CpGs located outside PMD boundaries experienced either a slight gain of methylation if they were located near other CpGs (‘social’), or a slight loss of methylation if they were isolated ‘solo’ CpGs (**Fig. 1c**). For CpGs within PMDs, the rate of hypomethylation appears influenced by immediate context, again with ‘solo’ CpGs losing methylation more rapidly than ‘social’ CpGs, and specifically with solo-WCGWs experiencing the most dramatic methylation loss, which is consistent with previous cross-sectional static characterizations in tissues.

We investigated whether PD-dependent PMD solo-WCGW hypomethylation occurs in different cellular contexts. We observed that across a range of primary human cell types from different developmental stages, the median methylation of PMD solo-WCGWs is tightly anticorrelated with PDs (**Fig. 1d**). The starting median methylation varies across cell lines, suggesting that the tissues from which these lines were derived have distinct replicative histories—an observation consistent with the variation in donor age and source tissue. Additionally, the rates of global PMD solo-WCGW methylation loss vary between cell types, perhaps reflecting different landscapes of CpG behavior. The pattern of methylation loss at individual PMD solo-WCGWs was reproducible between biological replicates (**Fig. 1e**).

### PMD solo-WCGW methylation loss is driven by proliferation-associated DNA replication

Elapsed time is linearly correlated with PDs until near-senescence for each primary cell line with a constant rate of cell division (**Suppl Fig. 1a**). As a result, methylation at PMD solo-WCGWs also correlates strongly with time (**Supplementary Fig. 1b**). Therefore, the serial passage by itself cannot distinguish between time-dependent loss of DNA methylation versus hypomethylation driven by cell division. To determine whether PMD solo-WCGW methylation loss is driven by cell division, or merely ensues with the passage of time, we cultured primary human fibroblasts with media containing decreasing concentrations of fetal bovine serum to impose different proliferation rates. We found that decreased rates of cell division by serum deprivation caused a dose-dependent reduction in DNA methylation loss, consistent with proliferation-associated loss of PMD solo-WCGW methylation (**Fig. 1f-h**). We have previously hypothesized that PMD solo-WCGW methylation loss is driven by incomplete maintenance methylation. Evidence from other groups has found that the solo-WCGW context is maintained inefficiently, although replication-uncoupled methylation was able to compensate somewhat, at least for a single cell cycle^34^. To test whether methylation loss is indeed driven by proliferation-associated DNA synthesis, we transiently treated several primary cell lines for 3 hours either with mitomycin C (MMC), a DNA replication inhibitor that can achieve full permanent cell cycle arrest, or with vehicle control, and maintained the cells for several weeks free of drug (**Fig. 1i-k**). The DNA synthesis-blocked, growth-arrested cells did not lose significant methylation at PMD solo-WCGWs despite the passage of time, whereas vehicle controls continued to progressively lose methylation as a function of cell divisions.

Taken together, these results present the first experimental evidence of a direct causal relationship between proliferation-associated DNA synthesis and PMD solo-WCGW hypomethylation.

### Factors driving CpG methylation trajectories in primary and immortalized cells

We investigated factors that could influence the varied rates of methylation loss among CpGs and between cell lines. Despite the similar profiles of median PMD solo-WCGW methylation loss, we observed subtle cell-type differences at individual CpGs (**Supplementary Fig. 2)**. Expression of genes encoding *de novo* and maintenance methylation machinery and TET enzymes was not markedly different between cell lines and across PDs (**Supplementary Fig. 3**).

PMD solo-WCGWs were grouped into major categories (**Fig. 2a, Supplementary Fig. 4**); those that remained stably methylated through replicative senescence, those that displayed variable methylation (>10% change), and those that were stably unmethylated. Primary cell lines from chronologically older individuals displayed a smaller stably methylated group, and larger stably unmethylated group (**Supplementary Fig. 4**). The variably methylated group was the largest for most cell lines, and was comprised overwhelmingly of CpGs that lost methylation, although a minor subset gained methylation. We further split the variably methylated group for primary fibroblast line AG06561 into quartiles of initial methylation levels to visualize the consistency of methylation loss across a spectrum of starting methylation (**Fig. 2a**, dark right panels).

**Figure 2.**
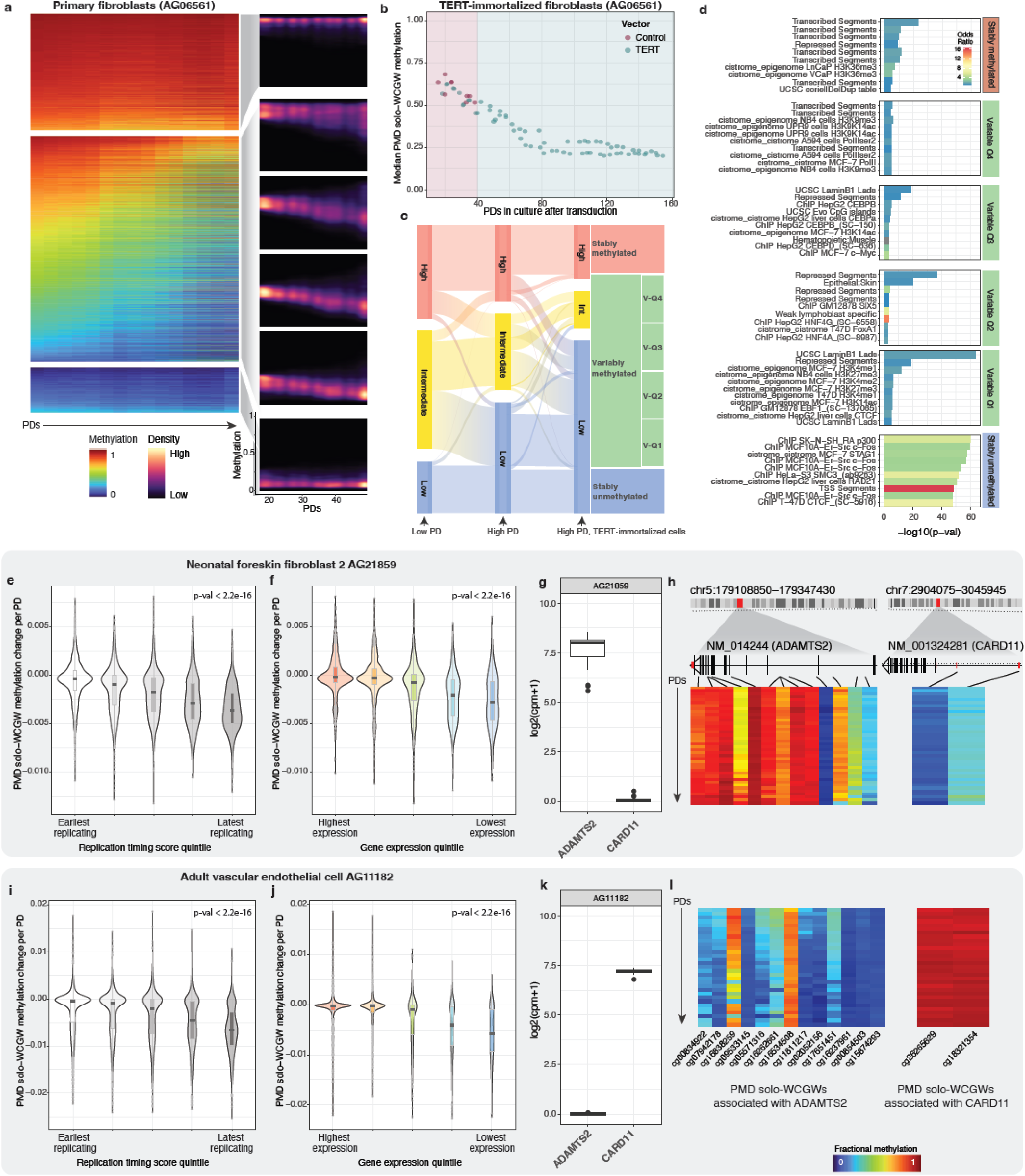
Meaningful groupwise PMD solo-WCGW behaviors. **a**, PMD solo-WCGWs were separated into major categories: (from top) stably hypermethylated, variable, and stably hypomethylated. Representative cell line AG06561 (fetal skin fibroblast) is depicted. Left, methylation heatmap of CpGs (rows) within each category. Samples (columns) are ordered by advancing population doublings (PDs). Right, density plot of probes within each major category, with the variable group further split into quartiles by starting methylation. **b**, Median PMD solo-WCGW methylation for TERT-immortalized and control primary fibroblasts through replicative senescence for control fibroblasts (pink shaded region) and through late PDs for immortalized fibroblasts (blue shaded region). **c**, Redistribution of PMD solo-WCGWs at early, non-immortalized passage, late, non-immortalized passage, and late, TERT-immortalized passage. **d**, PMD solo-WCGWs in TERT-immortalized fibroblasts were grouped by same paradigm in panel **a**. Locus overlap enrichment analysis was performed on each group, with all PMD solo-WCGWs on array as background. **e**, PMD solo-WCGW methylation change per population doubling (PD) for neonatal foreskin fibroblast 2 (AG21859) binned into quintiles based on ENCODE replication timing WA scores from BJ fibroblasts. **f**, methylation change per PD binned into expression quintiles of CpG-associated genes (primary RNA-seq data, AG21859). **g**, fibroblast gene expression for differentially expressed genes ADAMTS2 and CARD11. **h**, fibroblast DNA methylation heatmaps for PMD solo-WCGWs associated with differentially expressed genes ADAMTS2 (left) and CARD11 (right). Samples (rows) are arranged from early PD to late PD. **i**, PMD solo-WCGW methylation change per PD for adult vascular endothelial cell (AG11182) binned into quintiles based on ENCODE replication timing WA scores from HUVECs. **j**, methylation change per PD binned into expression quintiles of CpG-associated genes (primary RNA-seq data, AG11182). **k**, endothelial cell gene expression for differentially expressed genes ADAMTS2 and CARD11. **l**, endothelial cell DNA methylation heatmaps for PMD solo-WCGWs associated with differentially expressed genes ADAMTS2 and CARD11. Statistical comparisons for panels **e, f, I, j** by Kruskal-Wallis test.

To test whether there is a meaningful threshold of replicative history at which PMD solo-WCGW methylation stabilizes, primary fibroblasts (AG06561) were immortalized with a lentiviral construct carrying telomerase reverse transcriptase (TERT). DNA methylation was profiled at multiple passages following selection for both TERT-immortalized and control vector cells.

TERT-immortalized cells achieved drastically higher PDs than did control cells. We terminated the experiment after more than 150 PDs. At the last passage in this experiment, the TERT-immortalized cells remained highly proliferative (**Supplementary Fig. 5**). DNA methylation analysis indeed revealed a threshold at which PMD solo-WCGW methylation stabilized (**Fig. 2b**), approximately 40 PDs following replicative senescence of control cells. Although by the end of the experiment most CpGs had dropped to low levels of methylation, a small minority remained stably methylated (**Fig. 2c**).

We further investigated the distribution of residual methylation in high-PD TERT-immortalized cells. We used the methylation state to group CpGs into high, intermediate, and low methylation for early passage, late passage, and late TERT-immortalized cells (**Fig. 2c**). We identified CpGs that were stably methylated or stably unmethylated throughout, and split the remaining variably methylated CpGs into quartiles of terminal methylation values (**Fig. 2c**). The genomic coordinates of CpGs in each group were analyzed for enrichment of chromatin marks, genomic features, DNA binding proteins, and other characteristics that may explain their behavior (**Fig. 2d, Supplementary Table 2**). CpGs that were still highly methylated after extended post-immortalization culture had significant overlap with genomic features related to actively trasnscribed gene bodies. Among the top enriched overlapping features was H3K36me3, which is known to recruit *de novo* methyltransferase DNMT3B to transcribed gene bodies^37^. CpGs that had achieved low terminal methylation overlapped significantly with features bound by CTCF/cohesin complex members. The loss of methylation at sites bound by CTCF/cohesin complex members in severely hypomethylated immortalized cells is consistent with the chromosomal instability observed in severely hypomethylated model systems and human tumors^38–40^. We also observed an enrichment for hypomethylation at sites bound by c-Fos. We have previously shown that the AP-1 binding motif is overrepresented in genomic regions prone to hypomethylation in colorectal cancer^7^. We propose that DNA hypomethylation continues unabated upon TERT immortalization until finally reaching a severely hypomethylated equilibrium, in which compensatory *de novo* methylation offsets further demethylation. However, our bulk methylation analysis precludes investigating more population-driven explanations, such as selection against cells undergoing further loss of methylation.

### Replication timing and gene expression

To leverage our high-resolution methylation data into a more complete mechanistic understanding of PMD solo-WCGW dynamics, we regressed methylation to PDs at individual CpGs and compared the rate of methylation change to public replication timing annotations and primary gene expression data.

The short time window for maintenance re-methylation in late replicating regions is thought to contribute to hypomethylation at PMDs^6^. However, recent mechanistic studies indicate that maintenance methylation continues beyond S phase, uncoupled from the replication fork^34^. Although replication-uncoupled methylation mostly compensates for incomplete replication-coupled methylation following a single cell cycle^34^, its efficiency appears strongly influenced by neighboring CpG content, and the cumulative effect over many cell divisions has not been studied. PMD solo-WCGWs located in the regions replicating the latest lost methylation faster compared to those in earlier-replicating regions (**Fig. 2e,i**). This relationship suggests that PMD methylation loss is indeed driven by poor maintenance methylation, likely because of poor replication-coupled maintenance and subsequent failure of replication-uncoupled methylation. Other features co-occurring with late replication such as chromatin inaccessibility may further explain this relationship.

Enrichment analysis of CpGs that retained methylation at high PDs in TERT-immortalized fibroblasts (**Fig. 2d**) suggested that active transcription protects against replication-associated methylation loss. Although PMDs are relatively gene-poor^41^, there are several thousand gene-associated PMD solo-WCGWs on the EPIC array. Methylation change per PD was compared to expression level of associated genes. Indeed, high gene expression was protective against methylation loss (**Fig. 2f,j**). This relationship was cell-type specific; genes with differential expression between fibroblast line AG21859 and endothelial cell line AG11182 displayed alternate methylation at associated PMD solo-WCGWs (**Fig. 2g-h,k-l**). Genes with similar expression levels displayed similar methylation (**Supplementary Fig. 6**). We also examined whether the presence of H3K36me3 influenced the rate of methylation loss (**Supplementary Fig. 7**). Although there were few array PMD solo-WCGWs overlapping public annotations of this histone mark, its presence was significantly associated with reduced methylation loss for both cell types.

### Methylation loss during scheduled and unscheduled DNA synthesis

Culture characteristics are arguably non-physiologic; one with particular relevance to longevity research is oxygen exposure^42^. Chronic exposure to either high oxygen or reactive oxygen species results in premature aging phenotypes^43,44^. Primary cells grown in hypoxic chambers achieve more PDs^45–47^. To determine whether oxygen partial pressure affects PMD solo-WCGW dynamics in cultured cells, we serially cultured primary fibroblasts (AG21859) under ambient (approximately 20%) and low oxygen (3%) conditions (**Fig. 3a**), then performed DNA methylation profiling and RNA-sequencing across the series.

**Figure 3.**
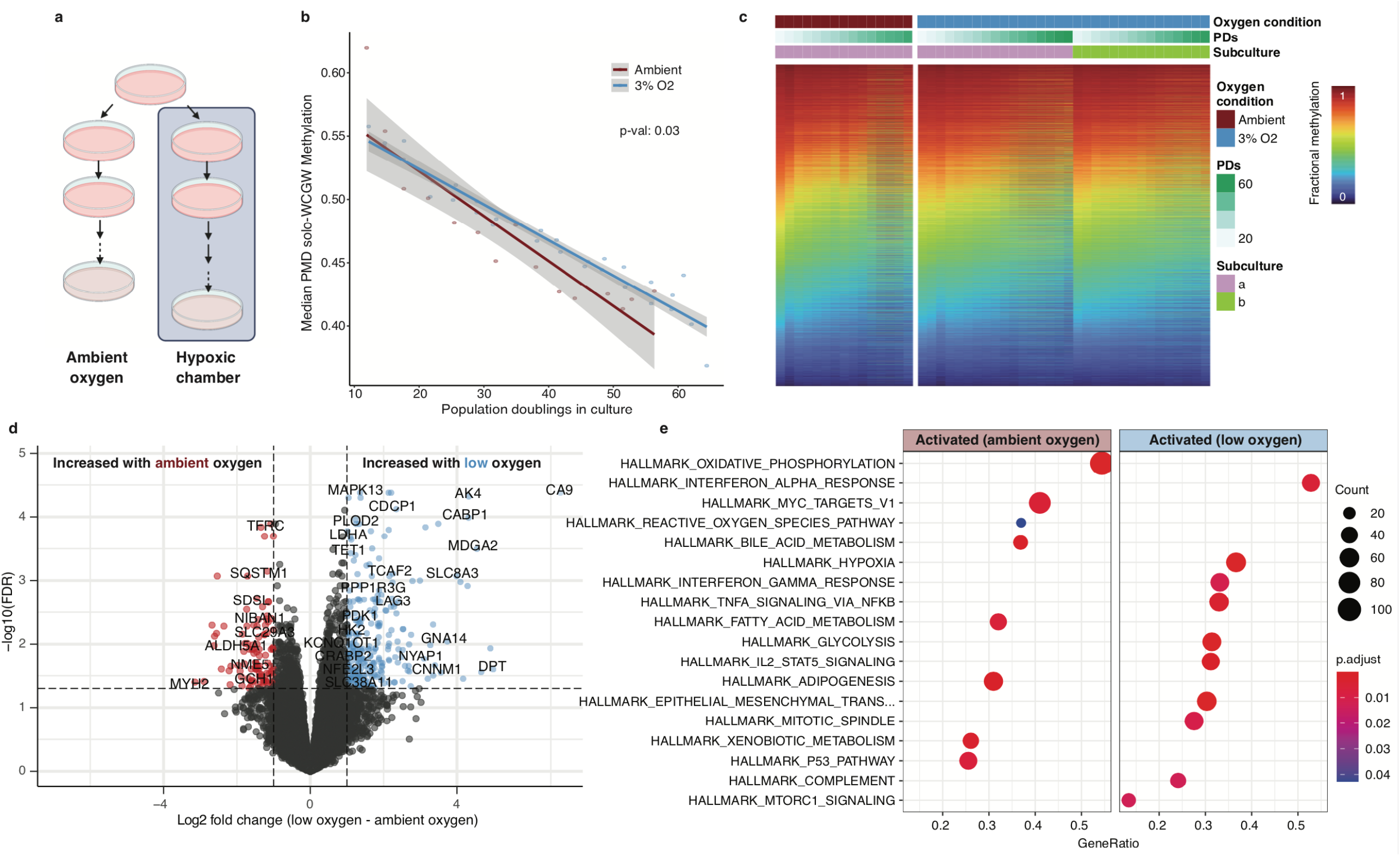
Low culture oxygen slows PMD solo-WCGW methylation loss. **a**, Schematic of tandem hypoxic/ambient oxygen primary cell culture. **b**, Median PMD solo-WCGW methylation plotted against population doublings (PDs) for both culture oxygen conditions. Solid lines depict linear regression with gray shading depicting 95% confidence interval. Statistical comparison of slopes by ANOVA. **c**, DNA methylation heatmap of PMD solo-WCGWs for both culture oxygen conditions. **d**, Volcano plot of differentially expressed genes between low oxygen and ambient oxygen culture. **e**, Pathway enrichment analysis results.

Primary cells grown under low oxygen conditions indeed achieved more PDs before replicative senescence than those grown under ambient oxygen conditions (**Supplementary Fig. 8**). Interestingly, median PMD solo-WCGW methylation loss was slowed under low oxygen culture (**Fig. 3b**). Individual CpGs behaved similarly across PDs between conditions (**Fig. 3c**), suggesting that cells grown in low oxygen conditions simply lose methylation more slowly.

Gene expression analysis between cells grown under both conditions revealed 641 genes significantly upregulated and 373 genes significantly downregulated under low oxygen culture (**Fig. 3d, Supplementary Table 3**). Top-upregulated genes in the low oxygen condition included many well-known hypoxia markers, such as carbonic anhydrase 9 (CA9) and adenylate kinase 4 (AK4), validating the experimental system and accompanying gene expression analysis. Top hits from differential pathway analysis included multiple metabolic pathways, pro-inflammatory pathways activated under low oxygen culture, and reactive oxygen species pathway activated under ambient oxygen culture (**Fig. 3e**).

While the bulk of DNA synthesis and accompanying DNA methylation maintenance occurs during the cell cycle^48,49^, a smaller amount also occurs during unscheduled DNA synthesis (UDS)^50^. UDS-coupled methylation maintenance efficiency is also sensitive to CpGs context^51^. We hypothesize that the accelerated methylation loss at PMD solo-WCGWs cultured in ambient oxygen is caused by incomplete methylation maintenance accompanying UDS as a consequence of increased oxidative damage^52^. This presents a minor caveat to using PMD solo-WCGW methylation as a proxy for replicative history. Conversely, the measure might also be useful to sensitively track cumulative oxidant/DNA damaging agent exposure in slowly proliferating cells or tissues.

### RepliTali: Modeling estimates of cumulative cell divisions

While median PMD solo-WCGW methylation correlates strongly with cell divisions in culture through standard replicative lifespans, we developed a more refined metric, which we named ‘RepliTali’ (for <u>Repli</u>cation <u>T</u>imes <u>A</u>ccumulated in <u>Li</u>fetime) to estimate relative replicative histories of human cells and tissues. PMD solo-WCGWs experience dramatic replication-associated methylation loss and are therefore depleted of methylation with relatively few cell divisions. To access a wider dynamic range of replication-associated methylation loss, we expanded the pool of eligible model CpGs to those in all sequence contexts within common PMDs. Since the total number of cell divisions prior to establishment of primary cell culture in our system is unknown, we envision this tool to be useful as a relative measure as opposed to an absolute benchmark for mitotic history. To adjust for variations in the in vivo replicative histories of the primary cells, we trained RepliTali upon normalized PDs (**Fig. 4a, Supplementary Table 4**).

**Figure 4.**
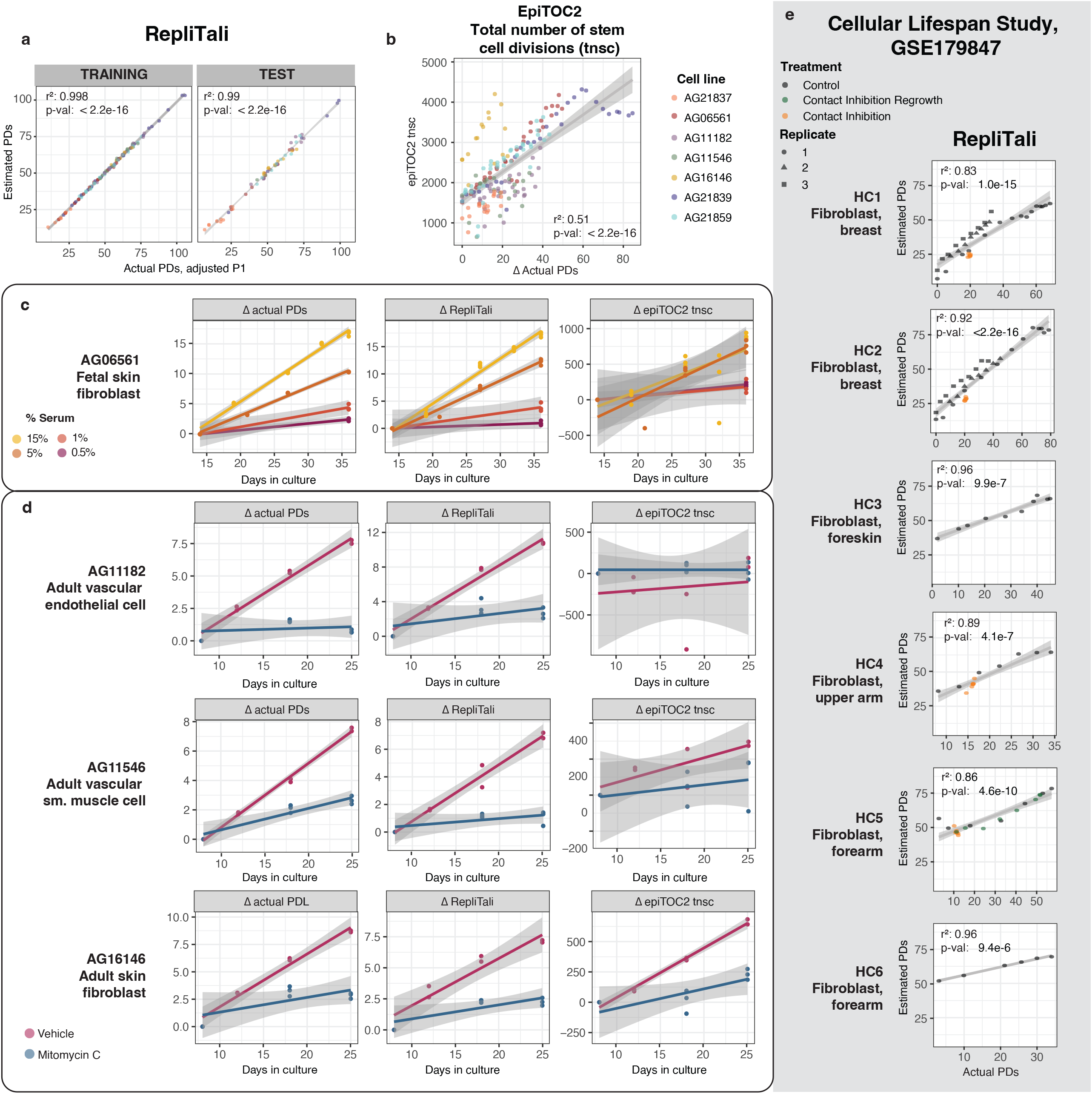
Construction and performance of RepliTali. **a**, Performance of RepliTali on randomized training (n=122) and test (n=60) sets. Solid lines depict linear regression with gray shading depicting 95% confidence interval. PDs: population doublings. **b**, Performance of epiTOC2, a hypermethylation-based mitotic clock, on primary cell culture DNA methylation data. **c**, Model performance on: Primary fibroblasts (AG06561) grown with different concentrations (%v/v) of media serum to achieve different proliferation rates, **d**, Mitomycin C (MMC) treated primary cell lines (n=3). **e**, Performance of RepliTali on external methylation dataset of cultured fibroblast lines (n=6).

### Comparing RepliTali performance to other models

We applied other published DNA methylation-based ‘mitotic clocks’^35,53–55^ to our primary cell data (**Fig. 4b, Supplementary Fig. 9a**). Performance of these models was not as tight and appeared highly cell line specific. Interestingly, hypermethylation-based clocks epiTOC2, pcgtAge, and MiAge were vulnerable to cell type differences, whereas epiCMIT^55^, a clock that selects the higher predicted mitotic age from either a set of CpGs that gains or a set that loses methylation, performed remarkably well on all cultured cell types. This is particularly interesting as epiCMIT was created exclusively from hematopoietic cell DNA methylation data.

Since the published ‘mitotic clocks’ were not trained on measured cell divisions, but rather on comparisons between different time points, it is important to investigate the extent to which RepliTali and these other clocks are reflecting time versus cell division. Our cell-cycle-attenuated and -arrested cell cultures are the best way to disentangle these two factors. Although RepliTali was trained on methylation data from primary cells cultured under standard conditions, it performed very well on growth-attenuated (**Fig. 4c**) and -arrested (**Fig. 4d**) primary cells, successfully distinguishing between divisions and time. Other existing ‘mitotic clocks’ performed inconsistently, with cell line-dependent performance again observed for hypermethylation-based clocks (**Fig. 4c-d, Supplementary Fig. 9b-c**).

### Validating RepliTali on External Datasets

We tested the performance of RepliTali and other clocks on a recent DNA methylation dataset of serially cultured primary fibroblasts^56^ (**Supplementary Table 5**). RepliTali performed strongly across all fibroblast lines, closely tracking with PDs under standard culture conditions (**Fig. 4e**). For cells growth-arrested via long term contact inhibition, RepliTali was very stable. Median PMD solo-WCGW methylation also performed well on this external dataset (**Supplementary Fig. 10**), supporting its use as a measure of replicative history, perhaps on non-EPIC array methylation datasets. Other mitotic clocks had varied performance, appearing sensitive to variations between fibroblast lines (**Supplementary Fig. 11)**. Again, epiCMIT outperformed exclusively hypermethylation-based clocks. Finally, we applied RepliTali and other mitotic clocks to several cell lines that have been extensively profiled (**Supplementary Fig. 12**). All clocks estimated colon adenocarcinoma-derived cell lines SW480 and HCT15 as having extremely high replicative histories. Curiously, the three hypermethylation-based clocks estimated that IMR90, a cell line initially derived from fetal lung fibroblasts that has been extensively cultured, had a replicative history comparable to low-passage primary skin fibroblast AG06561, whereas RepliTali and epiCMIT estimated higher values.

Whereas RepliTali was calibrated on actual, observed PDs accumulated in culture, other mitotic clocks were created using cancer data^54,55^ or normal aging blood^35,53^ data, with the assumption that malignant or aged tissues have experienced more cell divisions than non-malignant tissue. However, it is possible that CpGs prone to DNA methylation events co-occuring with, but not directly attributable to increased mitotic history in cancer and aging have been selected into these models. Additionally, past mitotic clocks were developed using the Infinium HumanMethylation450 (450K) array. This may explain why PMD solo-WCGWs have not yet been selected *en masse* as a tool for estimating mitotic age; they are severely underrepresented on this platform. Approximately 11% of genomic CpGs are PMD solo-WCGWs^33^, yet they comprise only 1.5% of 450K array CpGs. The relatively few (n=6214) on the 450K array were likely included because they overlap an enhancer or other gene regulatory feature, and thus often do not display the characteristic behavior of PMD solo-WCGWs. PMD solo-WCGWs represent approximately 27% of the probes in epiCMIT’s hypomethylation probeset, vastly exceeding the 1.5% represented on the 450K array. EpiCMIT had arguably stronger performance on our data and on the external dataset than the hypermethylation-based mitotic clocks. By comparison, while PMD solo-WCGWs comprise 18 of 87 CpGs in RepliTali, their mean coefficient weight was -3.35, versus a mean coefficient weight of -0.73 of all RepliTali CpGs, indicating that PMD solo-WCGWs contribute heavily to the model. Additionally, RepliTali CpGs recapitulated the progressive methylation loss behavior of PMDs at large (**Supplementary Fig. 13**).

Epigenetic clocks, representing models based on the methylation status at typically dozens to hundreds of CpGs, have become ubiquitous. Despite the astounding power of these models to predict features associated with biological aging—and its reversal^27^—the biological underpinnings of the CpGs that make these clocks ‘tick’ are often poorly understood. RepliTali is a new DNA methylation-based estimator of replicative history. Among methylation ‘clocks’ it is unique both in its construction—finely tuned upon serially passaged primary cells—and in our understanding of its driving mechanisms. RepliTali outperforms other models both on our own data and on an extensive external dataset. A challenge of developing a methylation clock to track mitoses is the highly variable rates of cell divisions between tissues^31^. However, the ability to dissect replicative history from other aspects of biological aging (perhaps simultaneously measured by another methylation clock) will aid our understanding of the aging process and inform therapies that seek to slow or reverse it.

## METHODS

### Primary Cell culture

All primary cell lines were obtained from the NIA Aging Cell Culture Repository at the Coriell Institute for Medical Research and cultured under recommended conditions. Fetal skin fibroblast AG06561 was maintained in Eagle’s MEM with Earle’s salts and non-essential amino acids (Gibco 11140-050) with 15% v/v fetal bovine serum. Neonatal foreskin fibroblasts AG21859 and AG21839 were maintained in Ham’s F12/DMEM 1:1 media supplemented with 10% v/v fetal bovine serum. Neonatal foreskin keratinocyte AG21837 was maintained in serum-free human epidermal keratinocyte media (MilliporeSigma SCMK001) on collagen IV-coated dishes (Corning 354453). Adult skin fibroblast AG16146 was maintained in Eagle’s MEM with Earle’s salts with 10% v/v fetal bovine serum. Vascular endothelial cell AG11182 was maintained in Medium 199 with 1X GlutaMAX (ThermoFisher 35050061), 0.02mg/ml endothelial cell growth supplement (Corning 354006), 0.05 mg/ml sodium heparin (Alfa Aesar A16198MD) and 15% v/v fetal bovine serum on plates pre-coated with gelatin (MilliporeSigma ES006B). Vascular smooth muscle cell AG11546 was maintained under the same conditions as AG11182 with the exception of 10% v/v fetal bovine serum. All primary cells were maintained at 37°C and 5% CO2 with ambient O2 unless otherwise noted.

Triplicate cultures derived from the same parent plate or vial obtained from Coriell were maintained in parallel through replicative senescence, which was defined in this study as drastically slowed growth (inability to reach near-confluence at 14 days after previous passage) or viable fraction of cells falling below 60%.

Passaging occurred as cells became approximately 90% confluent. At each passage, one fraction of cells was pelleted and frozen for future nucleic acid extraction. Another fraction was kept in suspension at room temperature and counted on an automated hemocytometer (BioRad TC20) in duplicate. Viability was determined by trypan blue dye exclusion.

Cumulative cell divisions in culture (population doublings, PDs) were determined using the following equation:

PD = 3.32 (log(cell yield)—log(viable cell inoculum) + X, with X being the PD of the inoculum

### Mitomycin C Treatment

Primary cell lines AG11182, AG11546, and AG16146 were reintroduced into culture from cryopreserved early-passage cells. Duplicate subcultures were derived from the initial recovered plate for treatment and control conditions. Cells were treated with DNA intercalating agent Mitomycin C (MMC, Alfa Aesar J63193MA) reconstituted in DMSO at a final concentration of 10ug/ml. An equal volume of DMSO was added to control subcultures. Both conditions were incubated for 3 hours at 37°C before media containing MMC or vehicle was removed, cells rinsed with PBS, and basal media replaced. Control cells were passaged normally, and growth-arrested, MMC-treated cells were collected on days 18 and 25.

### Primary Cell Growth Slowing

Primary fibroblast AG06561 was reintroduced into culture from cryopreserved early-passage cells. Four parallel cultures were established and were maintained in media containing 15%, 5%, 1%, and 0.5% v/v fetal bovine serum to encourage different rates of proliferation. At each passaging a fraction of cells was retained for DNA methylation analysis.

### TERT-immortalization

Low-PD primary fibroblasts (AG06561) were transduced with purified lentiviral particles containing expression vectors encoding human Telomerase Reverse Transcriptase (TERT) and hygromycin resistance marker (AMSBIO LVP1131-Hygro-PBS), or hygromycin resistance marker alone (control, AMSBIO EF1a-Null-Hygro). Following selection with 250ug/ml hygromycin B, cells were serially cultured either through replicative senescence (control) or in perpetuity (TERT). At the time of analysis, nearly a full year after transduction, the immortalized cells remained highly proliferative.

### Low Oxygen Cell Culture

Low-PD primary fibroblasts (AG21859) were cultured in triplicate in either a standard incubator (Panasonic MCO-19AICUVPA) with ambient O_2_, or dual-gas CO_2_/N_2_ incubator (PHCbi MCO-170M-PA) at 3% O_2_, through replicative senescence.

### Methylation Analysis by Microarray

Briefly, frozen cell pellets were thawed and lysed using QIAshredder spin columns (Qiagen 79656). Genomic DNA was extracted from each sample using the AllPrep DNA/RNA Mini Kit (Qiagen 80204), then stored at -80°C before analysis. DNA was quantified by Qubit fluorimetry (Life Technologies). Approximately 500ng of genomic DNA was bisulfite converted using the Zymo EZ DNA methylation kit (Zymo Research D5004) then hybridized overnight on an Infinium MethylationEPIC BeadChip (Illumina), in which the genomic DNA molecules anneal to locus-specific DNA oligomers linked to individual bead types. Raw signal intensities were exported as .idat files, which were processed using the R package SeSAMe^57,58^. Of 386 DNA methylation samples run, 14 failed quality control and were excluded from further analysis, producing a final analytical sample count of 372. All DNA methylation data can be accessed through the Gene Expression Omnibus (GEO) accession GSE197471.

### Statistical Analysis

Analysis was performed in R software (version 4.1.1). For comparisons of effect of MMC growth arrest and serum-dependent growth slowing on PMD solo-WCGW methylation, mixed-effects modeling (R package ‘lme4’) was performed, using logit-transformed (m) methylation values. Multiple comparisons were performed via Tukey contrasts. For comparison of the effect of culture oxygen condition on rate of PMD solo-WCGW methylation models were compared via ANOVA, again using logit-transformed (m) methylation values.

### LOLA

Genomic coordinates (hg19) of PMD solo-WCGW probes of interest were subject to Locus Overlap Enrichment Analysis (LOLA) using R package ‘LOLA’^59^ and LOLACore (hg19) region set database, available here: https://databio.org/regiondb.

Coordinates of all PMD solo-WCGW probes on the InfiniumEPIC Methylation array were used as background for enrichment analysis.

### RNA-seq

RNA was isolated from frozen cell pellets using the AllPrep DNA/RNA Mini Kit (Qiagen 80204), then stored at -80°C before analysis. RNA Libraries were prepared from 100 ng of total RNA with the KAPA Stranded mRNA-Seq Kit (Kapa Biosystems KK8401). Indexed libraries were then pooled and 2×50 bp, paired-end sequencing was performed on an Illumina NovaSeq 6000 sequencer to a minimum read depth of 30M reads/library. Differential expression analysis was performed with standard edgeR and DESeq2 workflow. Senescence timepoints were excluded from differential expression and pathway enrichment analysis for oxygen culture condition experiment. Scripts for RNA-seq analytical workflow, including downstream analysis in R, are available here: https://github.com/vari-bbc/rnaseq_workflow

### RepliTali Construction

Starting PD values of primary cell lines were normalized using an elastic net regression model (R package ‘glmnet’) trained on the chronologically youngest cell line, fetal skin fibroblast AG06561. Samples from all cell lines were randomized into training (n=122) and test (n=60) sets; normalized PDs were used to construct the final ‘RepliTali’. RepliTali is constructed using array CpGs within common PMD boundaries. Coefficients are presented in **Supplementary Table 4**.

### Mitotic clock comparisons

EpiTOC estimates were obtained using the R script available at: https://zenodo.org/record/2632938#.YdWva5DMKrc. Script was run separately on each cell line, per the author’s specifications.

MiAge estimates were calculated with materials deposited here: http://www.columbia.edu/~sw2206/softwares.htm

epiCMIT estimates were calculated as described in https://duran-ferrerm.github.io/Pan-B-cell-methylome/Estimate.epiCMIT.html

### External Data

InfiniumEPIC Methylation probe manifest^60^ is available here: https://zwdzwd.github.io/InfiniumAnnotation

Common PMD coordinates, as well as coordinates and characteristics of PMD solo-WCGWs genome-wide and present on the InfiniumEPIC Methylation array are documented here:

https://zwdzwd.github.io/pmd

### Replication timing

Replication timing data from BJ foreskin fibroblasts and HUVECs was generated by the University of Washington and maintained by ENCODE. Files are available here: http://genome.ucsc.edu/cgi-bin/hgFileUi?db=hg19&g=wgEncodeUwRepliSeq Replication timing weighted average (WA) scores were calculated as previously specified^61^:

WA=(0.917*G1b)+(0.750*S1)+(0.583*S2)+(0.417*S3)+(0.250*S4)+(0*G2)

### H3K36me3

Histone ChIP-seq data from neonatal foreskin fibroblasts was generated by Joseph Costello’s lab at UCSF/Roadmap Epigenomics Project. Histone ChIP-seq data from HUVECs was generated by the University of Washington/ENCODE project. Neonatal foreskin fibroblast: ENCSR889OUV | GSM817238 HUVEC: ENCSR000DVM | GSM945233

### Methylation data

Infinium MethylationEPIC array data from serially passaged human fibroblasts was generated by Martin Picard’s lab at Colombia University (Cellular Lifespan Study 1.0^56^, GSE179847). Raw idats were reprocessed as above.

## ACKNOWLEDGEMENTS

This work was supported by NIH/NIA grant R01AG066764 awarded to PWL and HS, and by Van Andel Institute. The authors thank Dr. Wanding Zhou and Dr. Benjamin Berman for valuable input. The authors thank the Van Andel Genomics Core for providing DNA methylation array and RNA-seq facilities and services.

**Supplementary Figure 1.**
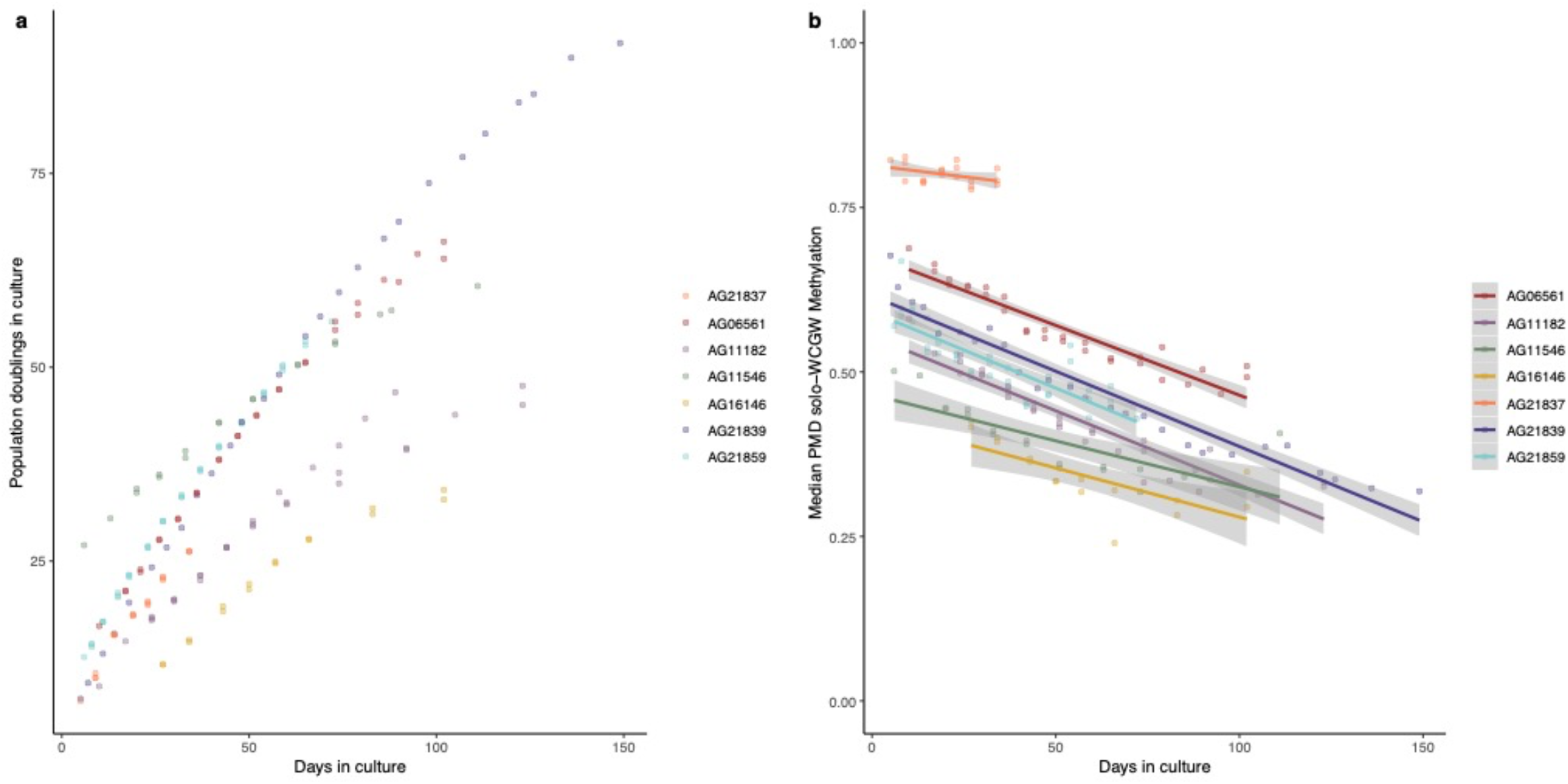
Correlation of median PMD solo-WCGW methylation with time. **a**, Growth curves of cell lines used in this study. **b**, Plot of median fractional methylation at 26,732 PMD solo-WCGWs present on InfiniumEPIC Methylation array vs. days in culture during this study. Regression lines are linear with shaded regions representing 95% confidence interval.

**Supplementary Figure 2.**
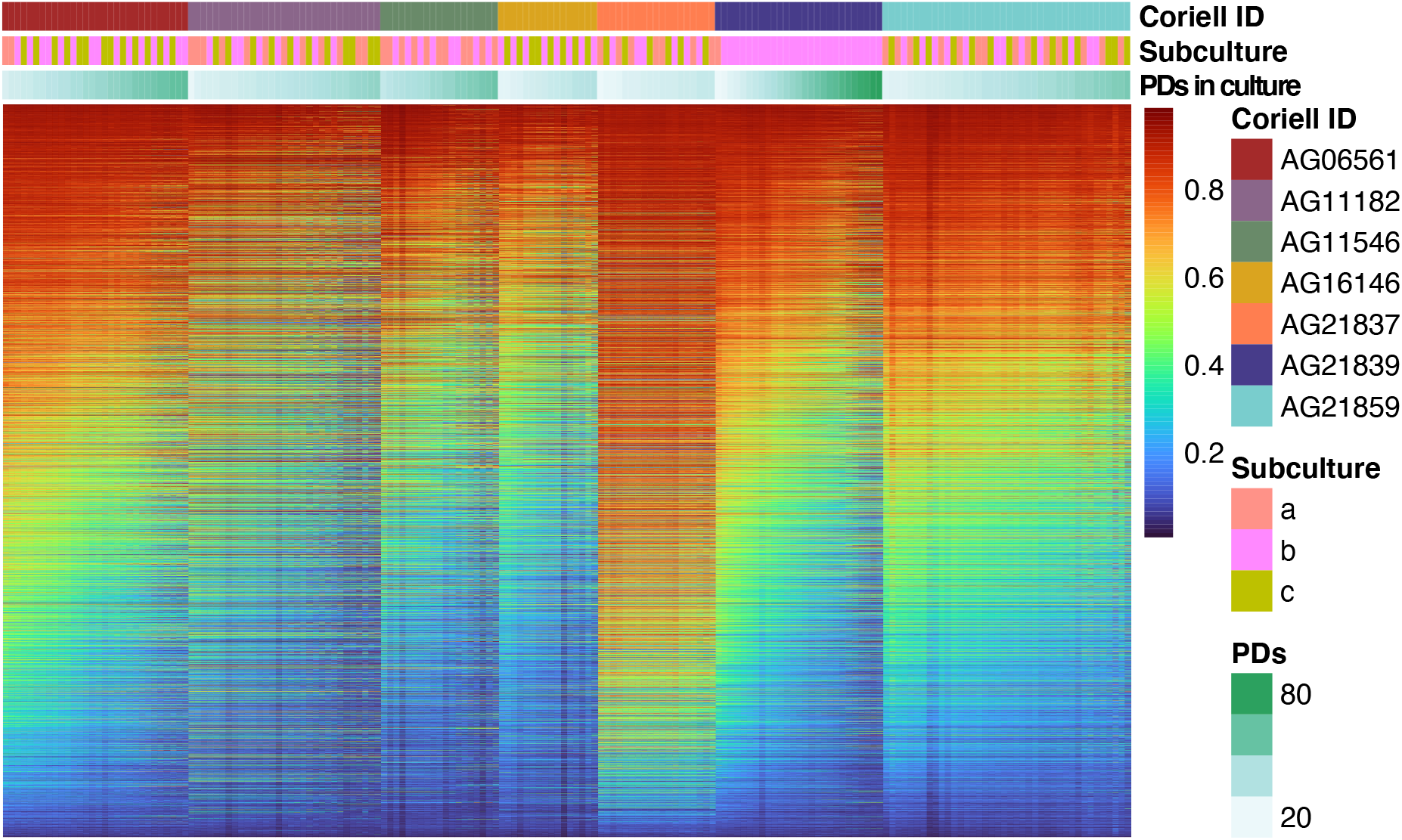
Heatmap of PMD solo-WCGW methylation for primary cell lines included in this study. Rows (CpGs) are ordered by the mean of each cell line’s median methylation value across timepoints profiled. Columns (samples) are ordered by advancing population doublings (PDs).

**Supplementary Figure 3.**
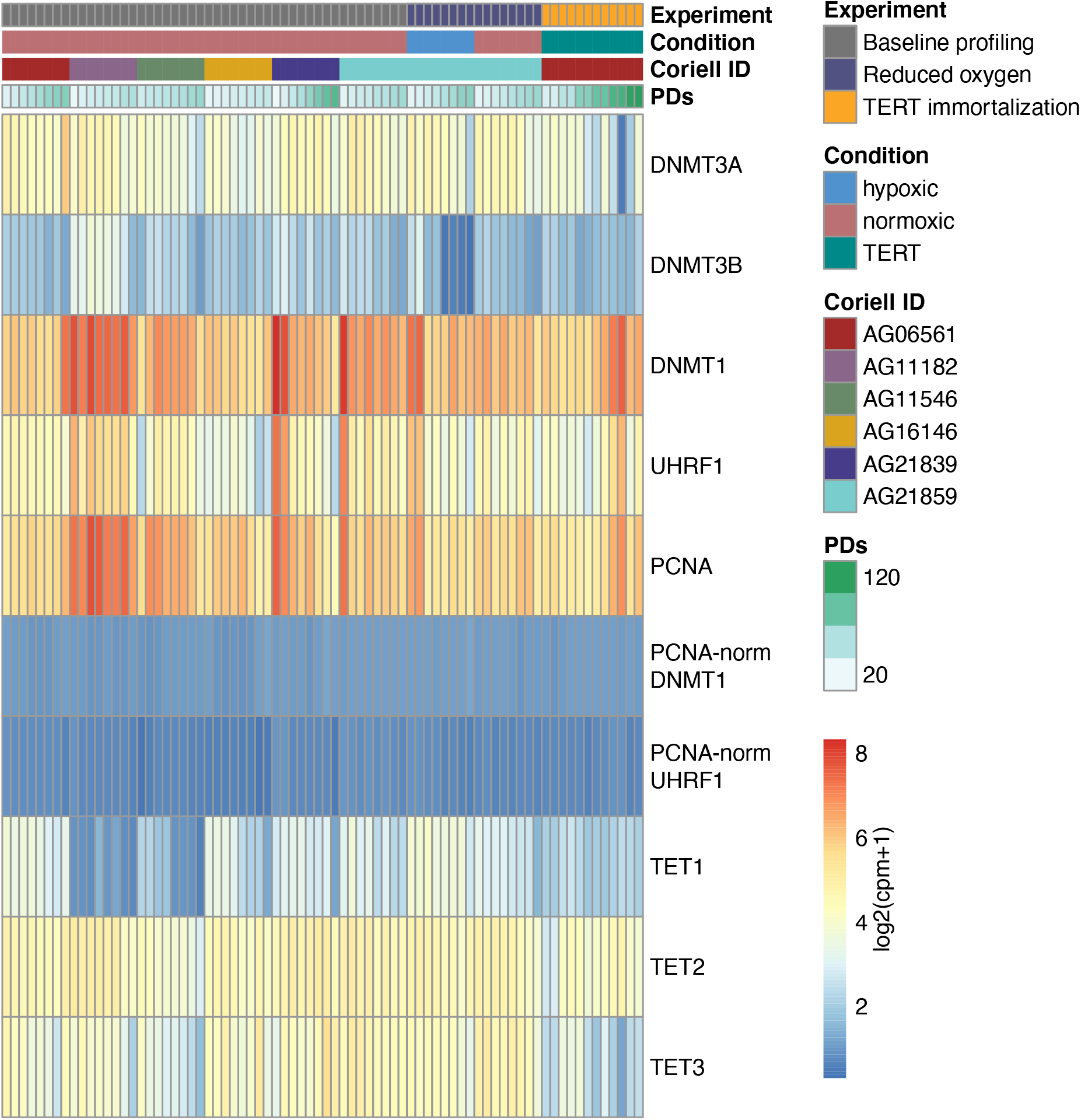
Gene expression heatmap of key enzymes involved in DNA methylation patterning. Columns (samples) are ordered first by experiment, then cell line, then by advancing population doublings (PDs).

**Supplementary Figure 4.**
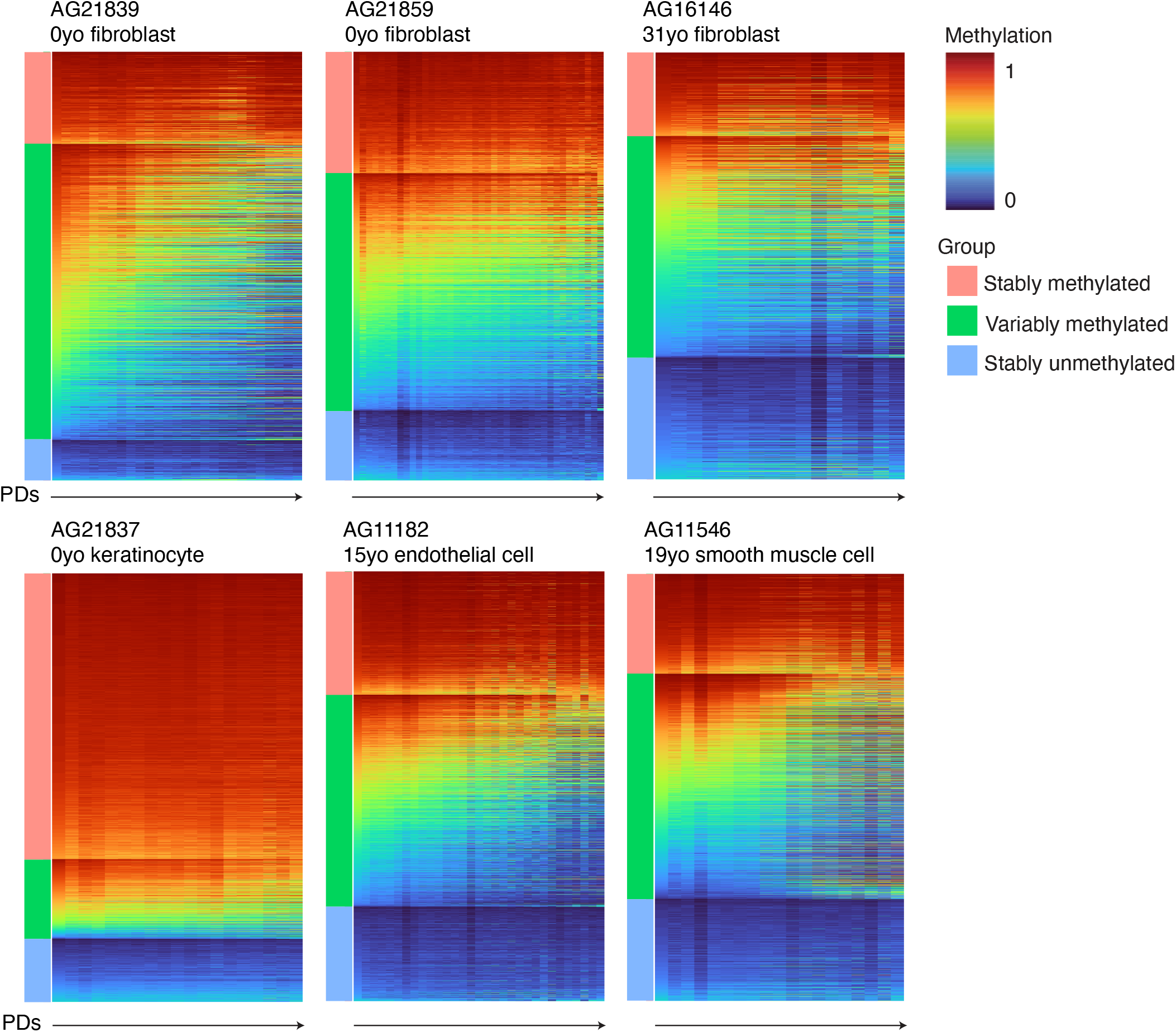
Methylation heatmaps of PMD solo-WCGWs. CpGs were separated into major categories: (from top) stably methylated, variably methylated, and stably unmethylated. Samples (columns) are ordered by advancing PDs.

**Supplementary Figure 5.**
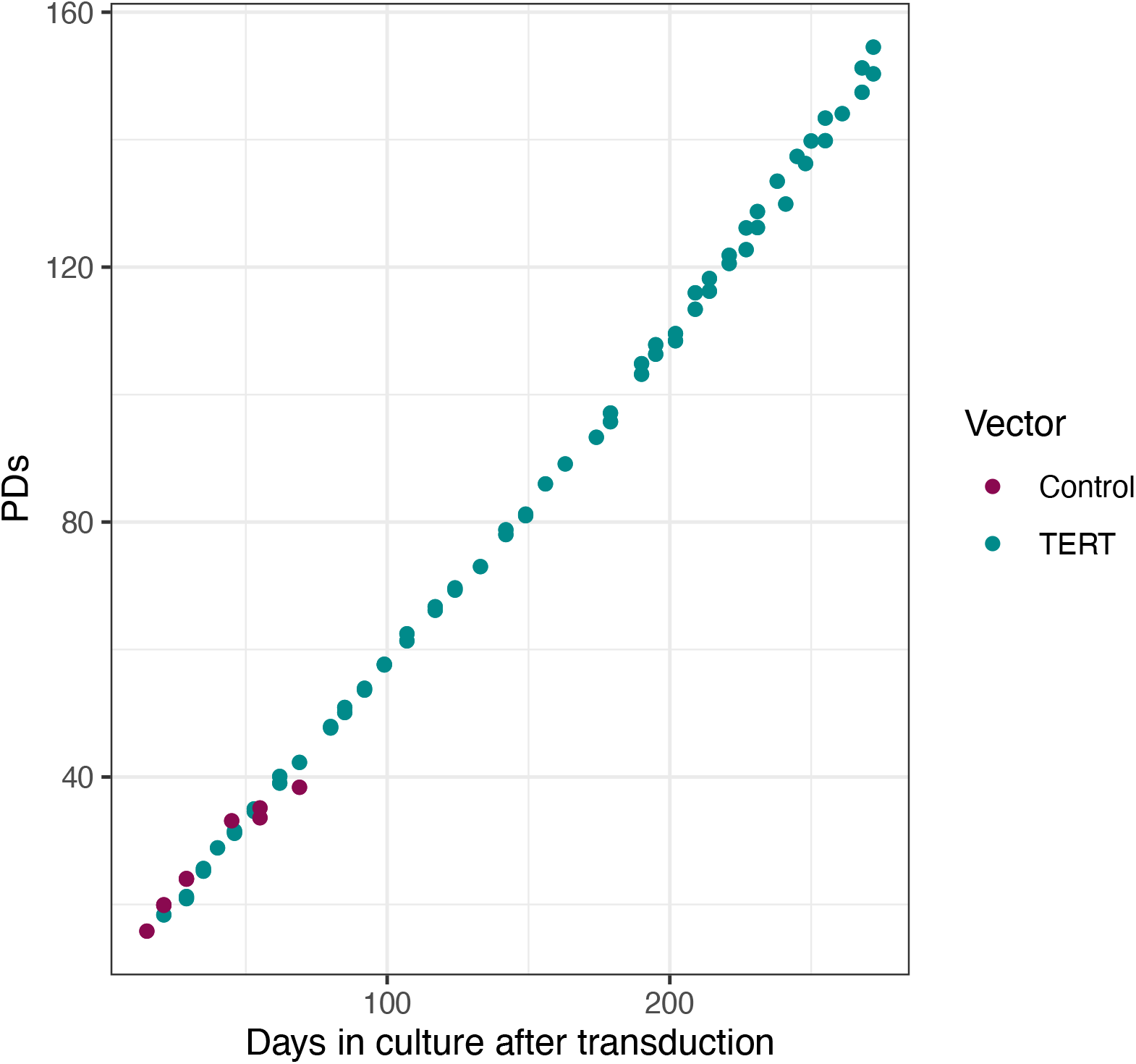
Growth curve of TERT-immortalized primary skin fibroblasts (AG06561). Control vector fibroblasts (dark pink) senesced at approximately 40 PDs; at the time of analysis TERT-immortalized cells (blue) remained highly proliferative.

**Supplementary Figure 6.**
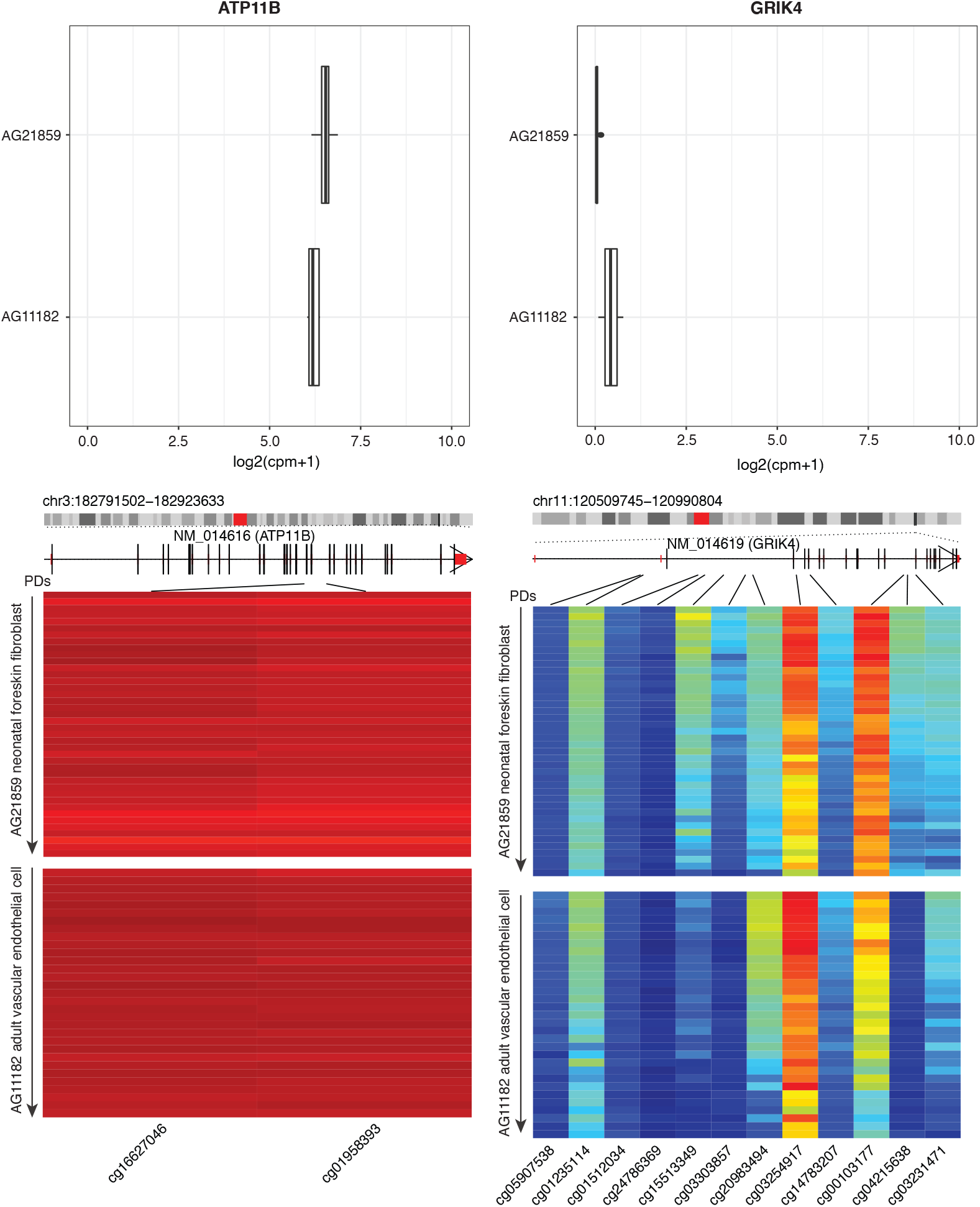
Gene expression protects against replication-associated methylation loss. Upper panels: boxplots of normalized gene expression for similarly expressed genes: ATP11B (left) and GRIK4 (right). Lower panels: methylation at gene-associated PMD solo-WCGWs. Samples (rows) are arranged from early PD to late PD.

**Supplementary Figure 7.**
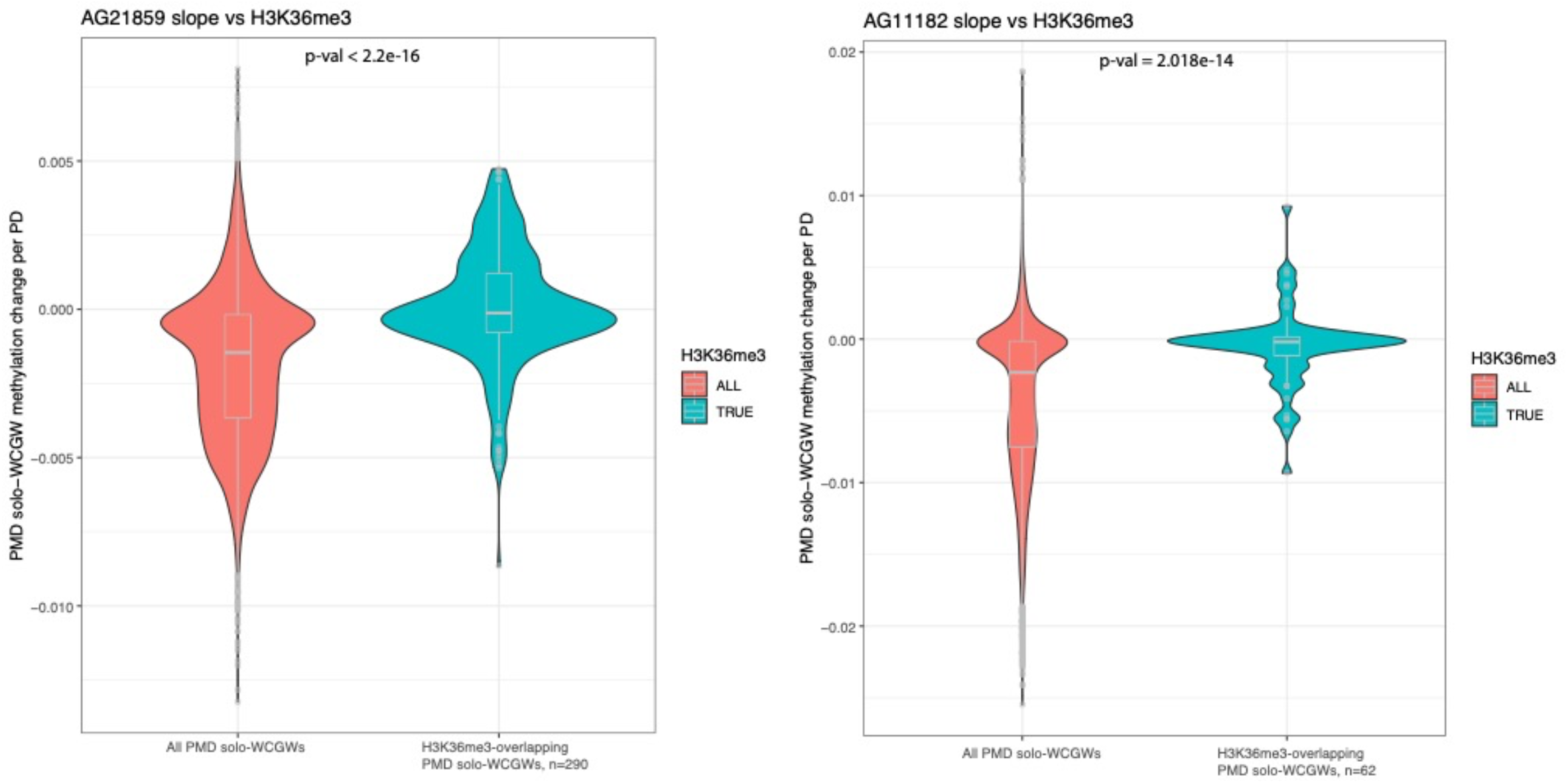
H3K36me3 protects against replication-associated methylation loss. CpG-wise methylation change per PD for PMD solo-WCGWs overlapping H3K36me3 for primary fibroblasts (AG21859, left) and endothelial cell (AG11182, right). Statistical comparison by Kruskal-Wallis test.

**Supplementary Figure 8.**
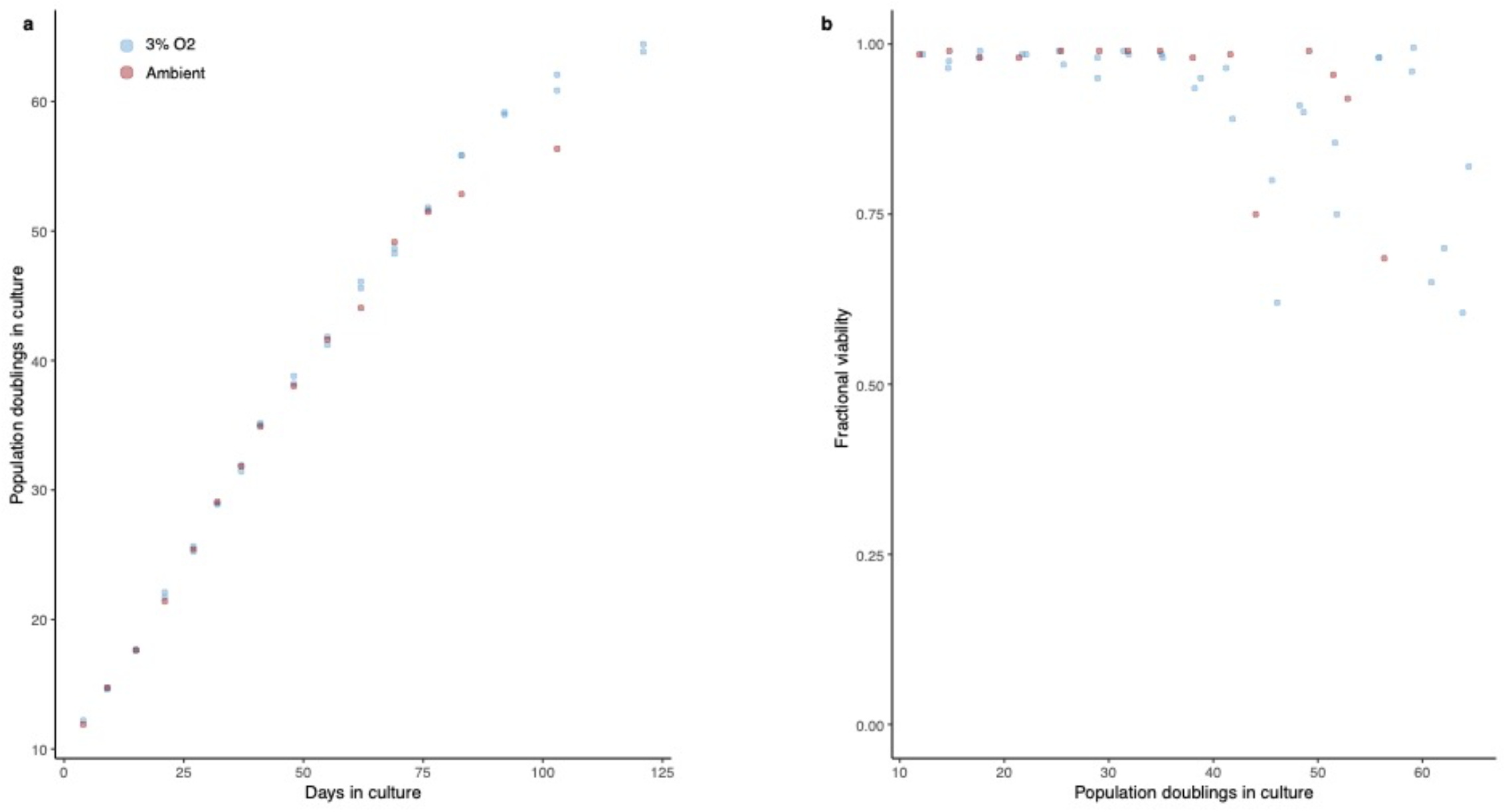
**a**, Growth curves of primary fibroblast (AG21859) grown under ambient oxygen conditions (pink) and low 3% oxygen (blue). **b**, Cellular viability of primary fibroblasts under both oxygen culture conditions.

**Supplementary Figure 9.**
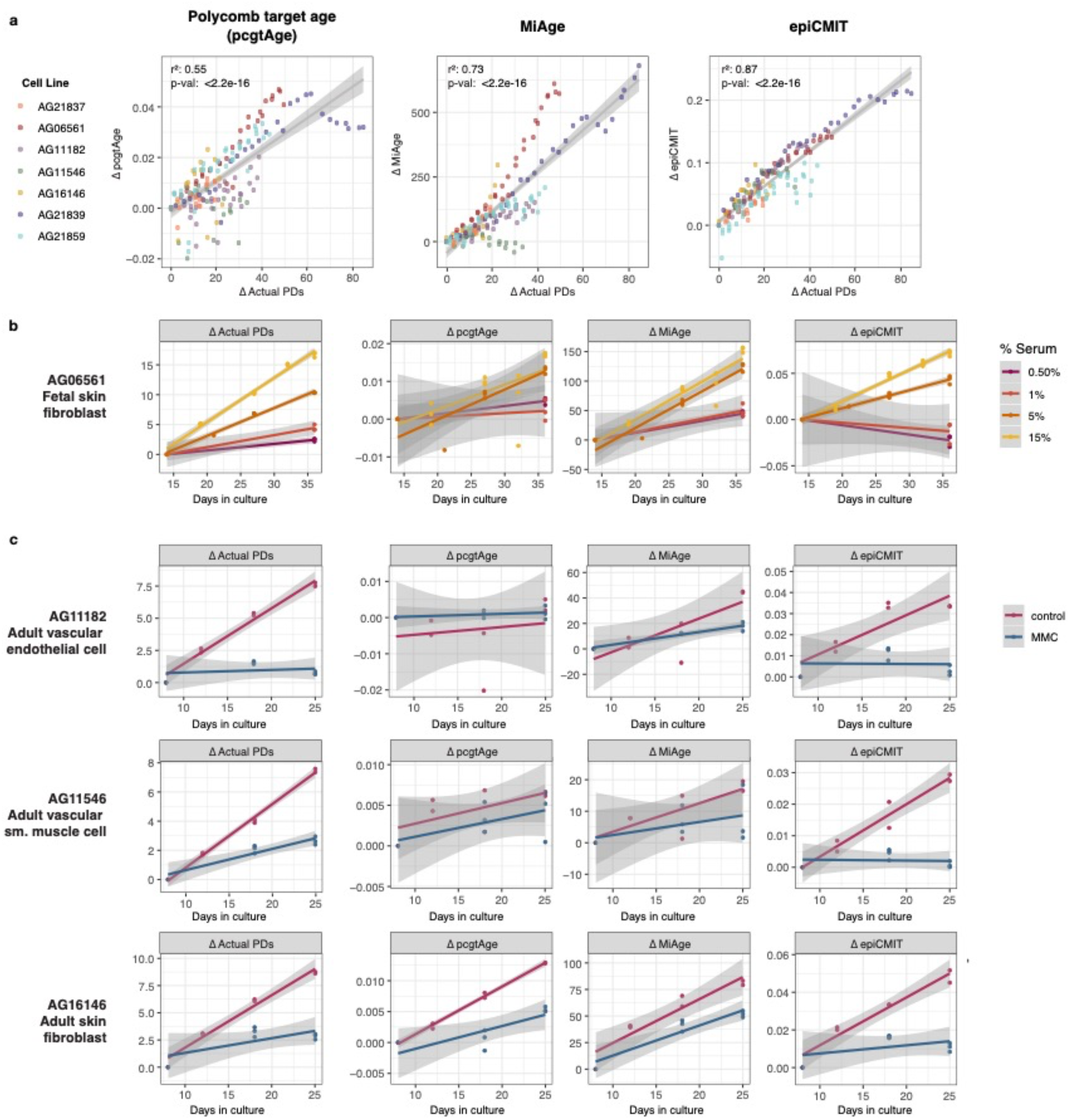
Performance of other mitotic clocks on primary DNA methylation data. Clock performance on (**a**) primary human cells cultured under standard conditions through replicative senescence; (**b**) primary human fibroblasts grown under different % v/v media serum to encourage differential proliferation rates; (**c**) primary human cells treated with DNA replication inhibitor Mitomycin C (MMC) or vehicle control.

**Supplementary Figure 10.**
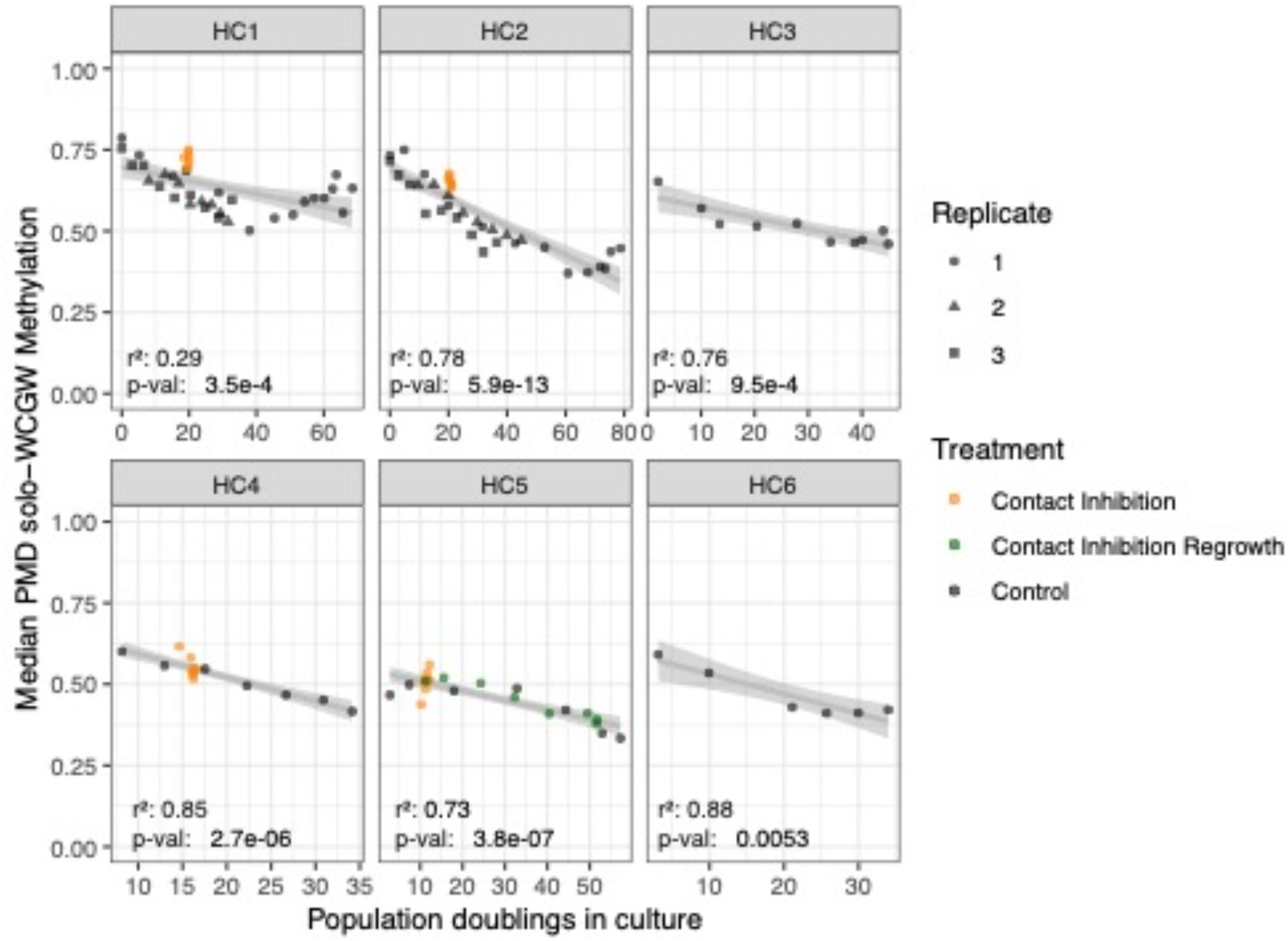
Loss of PMD solo-WCGW methylation as a consequence of cellular mitotic history, external DNA methylation dataset GSE179847.

**Supplementary Figure 11.**
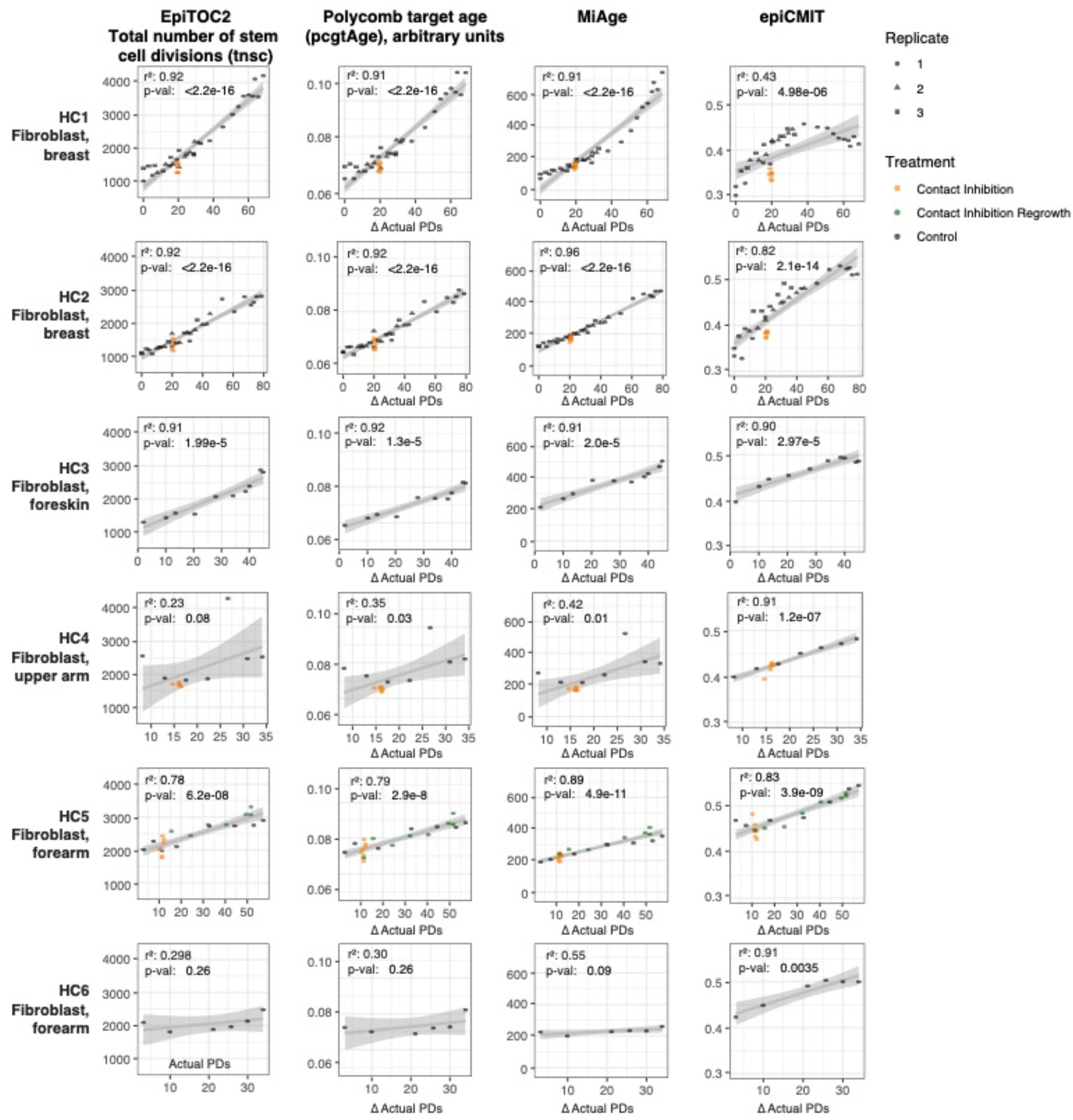
Performance of other mitotic clocks on external DNA methylation dataset GSE179847.

**Supplementary Figure 12.**
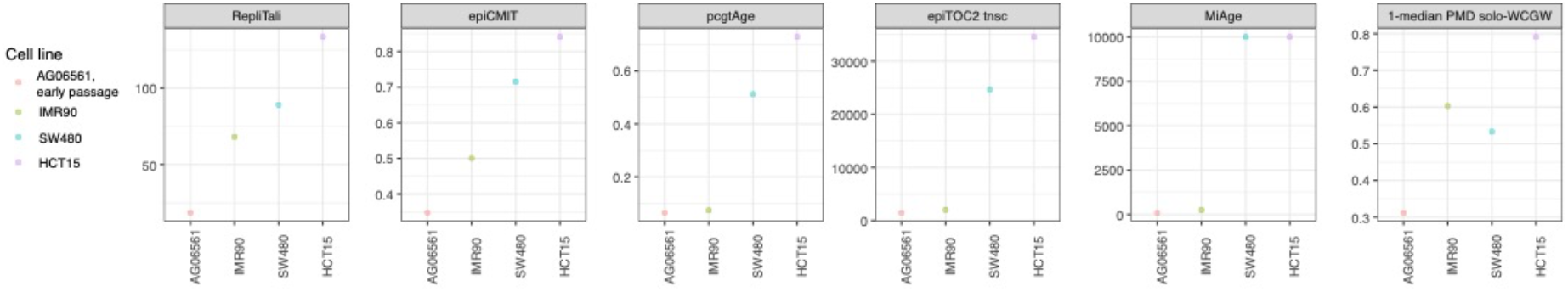
Mitotic clock estimates of replicative age on well-classified human cell lines. AG06561: Primary human fibroblast, early passage, for comparison. **IMR90**: Fetal lung fibroblast-derived cell line. **SW480**: Colon adenocarcinoma-derived cell line. **HCT15**: Colon adenocarcinoma-derived cell line.

**Supplementary Figure 13.**
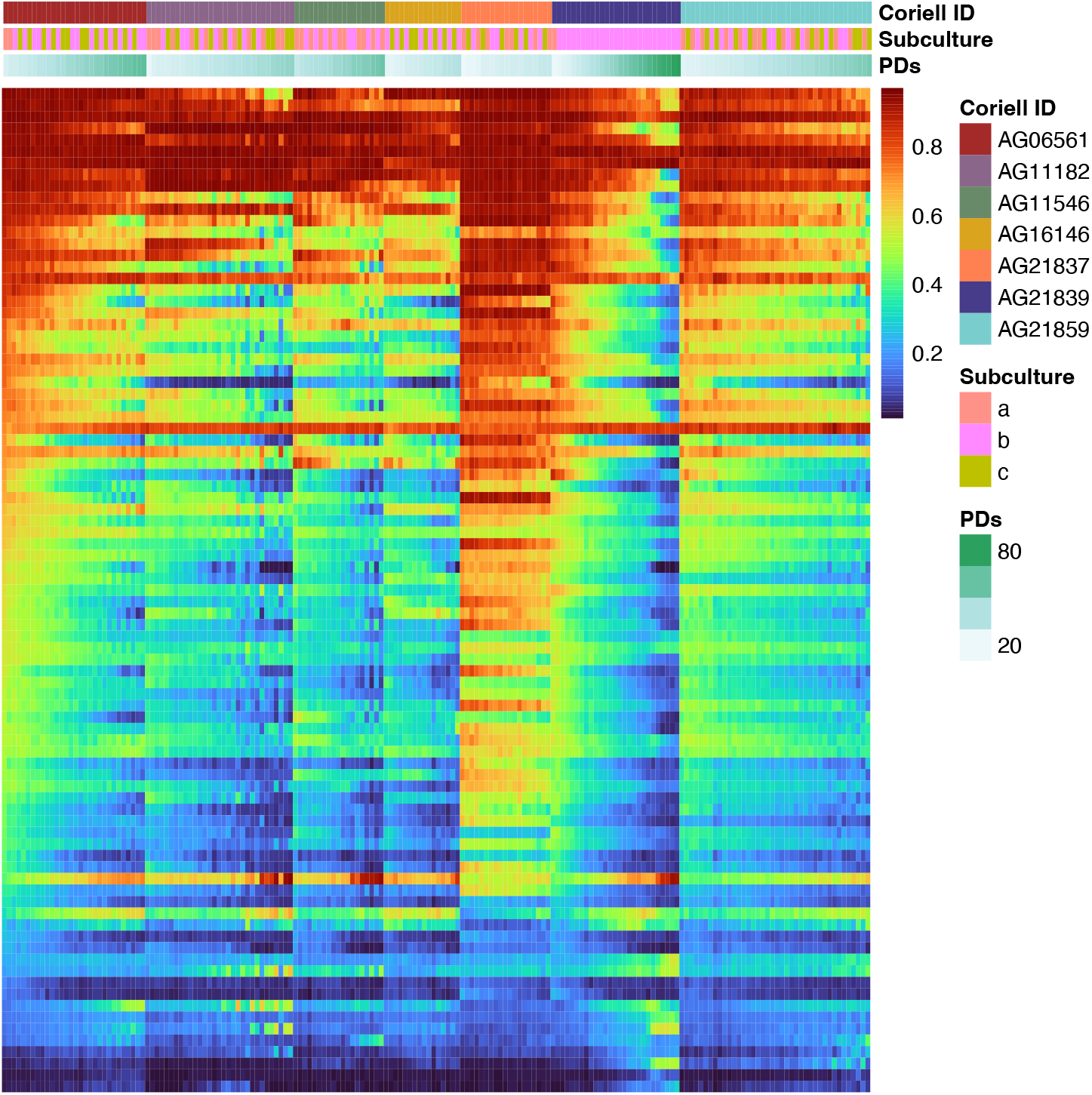
RepliTali CpG methylation for serially passaged cells.

**Supplemental Figure 14:**
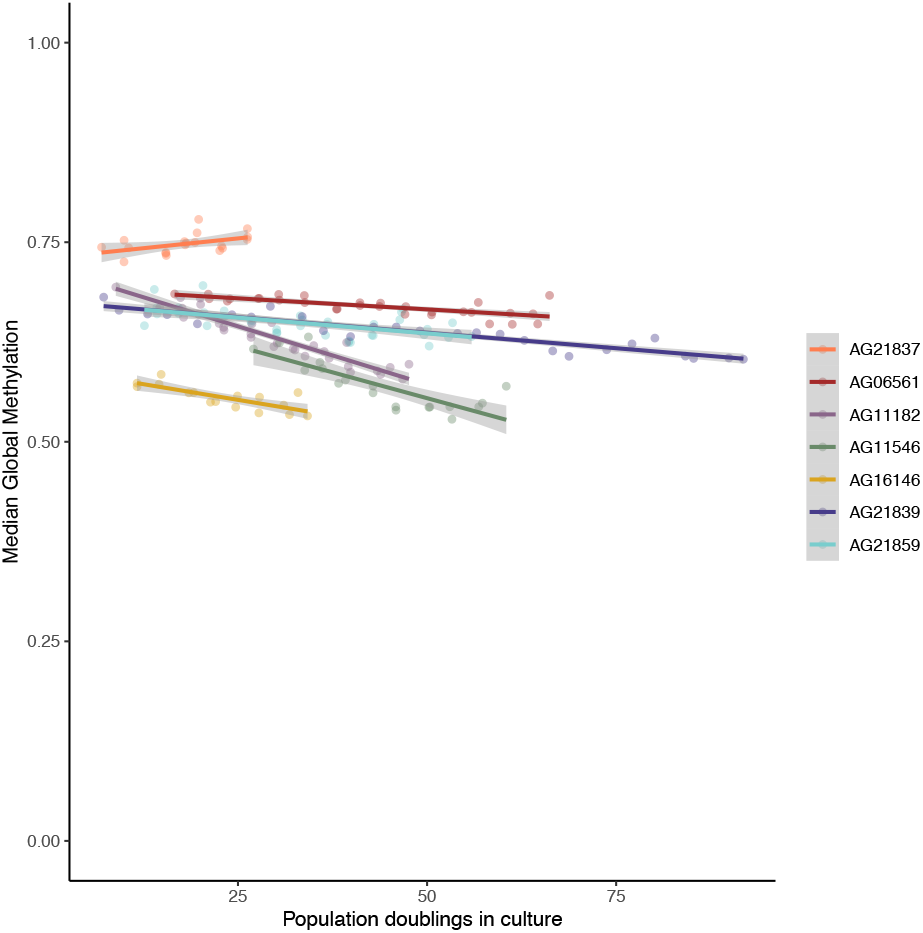
Global methylation in serially passaged primary cells grown under standard culture conditions.

**Supplemental Figure 15:**
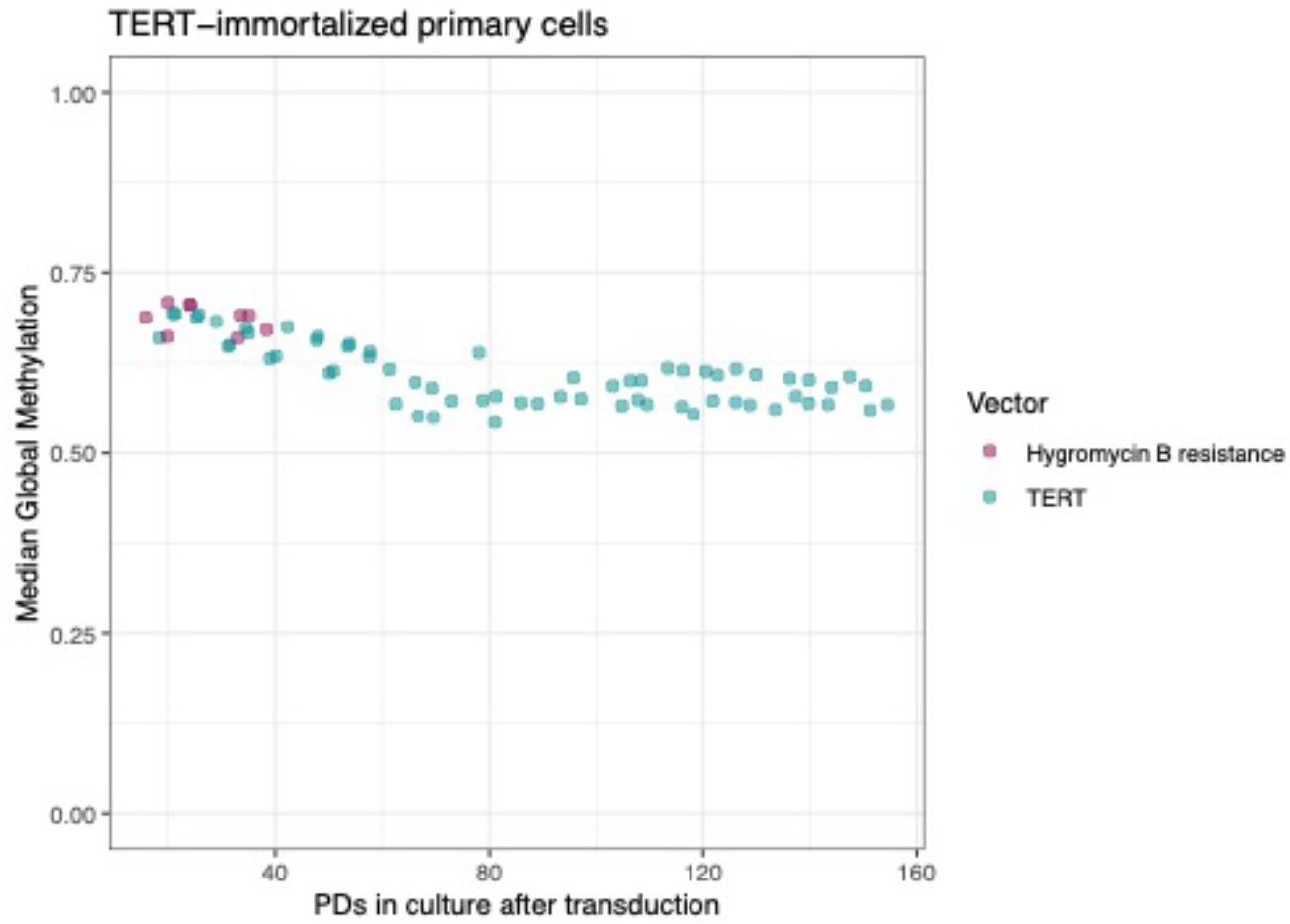
Global DNA methylation in immortalized primary fibroblast AG06561.

## Contents of additional materials

**Supplementary Table 1**: Summary of primary cell cultures used in this study.

**Supplementary Table 2**: Differential gene expression analysis for low oxygen vs ambient oxygen culture conditions

**Supplementary Table 3**: Groupwise Locus Overlap Enrichment Analysis results for TERT-immortalized fibroblasts

**Supplementary Table 4**: RepliTali coefficients

**Supplementary Table 5**: GSE179847 characteristics

