## Supplementary Table 1 for "Cell division drives DNA methylation loss in late-replicating domains in primary human cells"

**Endicott et al. Supplementary Material**

| **Cell line** | **Cell type** | **Donor tissue** | **Donor age** | **Sex** | **Documented starting**  **PDs (Coriell)** |
| --- | --- | --- | --- | --- | --- |
| AG06561 | Skin fibroblast | Skin, sacrum | 16fw | F | 12 |
| AG21859 | Skin fibroblast | Foreskin | 0y | M | 9.38 |
| AG21839 | Skin fibroblast | Foreskin | 0y | M | 5.39 |
| AG21837 | Keratinocyte | Foreskin | 0y | M | 2.33 |
| AG11182 | Endothelial cell | Vein, iliac | 15y | M | 22.9 |
| AG11546 | Smooth muscle | Vein, iliac | 19y | M | 26 |
| AG16146 | Skin fibroblast | Skin, arm | 31y | M | 9.7 |

**Supplementary Table 1**

Summary of primary cell cultures used in this study. Cell lines used were obtained from the NIA Apparently Healthy Collection at the Coriell Institute for Medical Research: <https://www.coriell.org/0/Sections/Collections/NIA/ApparentlyHealthy.aspx?PgId=324&coll=AG>
