## Supplementary Table 5 for "Cell division drives DNA methylation loss in late-replicating domains in primary human cells"

**Endicott et al. Supplementary Material**

| **Cell line** | **Cell type** | **Donor tissue** | **Donor age** | **Sex** |
| --- | --- | --- | --- | --- |
| HC1 | Fibroblast | Breast | 18y | M |
| HC2 | Fibroblast | Breast | 18y | F |
| HC3 | Fibroblast | Foreskin | 0y | M |
| HC4 | Fibroblast | Upper arm | 3y | M |
| HC5 | Fibroblast | Forearm | 29y | M |
| HC6 | Fibroblast | Forearm | 36y | F |

**Supplementary Table 5**

Characteristics of primary cell cultures in external methylation dataset GSE179847.
